# OligoTRAFTACs: A Generalizable Method for Transcription Factor Degradation

**DOI:** 10.1101/2021.12.20.473482

**Authors:** Kusal T. G. Samarasinghe, Elvira An, Miriam A. Genuth, Ling Chu, Scott A. Holley, Craig M. Crews

## Abstract

Dysregulated transcription factors (TFs) that rewire gene expression circuitry are frequently identified as key players in disease. Although several TFs have been drugged with small molecules, the majority of oncogenic TFs are not currently pharmaceutically tractable due to their paucity of ligandable pockets. The first generation of transcription factor targeting chimeras (TRAFTACs) was developed to target TFs for proteasomal degradation by exploiting their DNA binding ability. In the current study, we have developed the second generation TRAFTACs (“oligoTRAFTACs”) comprised of a TF-binding oligonucleotide and an E3 ligase-recruiting ligand. Herein, we demonstrate the development of oligoTRAFTACs to induce the degradation of two oncogenic TFs, c-Myc and brachyury. In addition, we show that brachyury can be successfully degraded by oligoTRAFTACs in chordoma cell lines. Furthermore, zebrafish experiments demonstrate *in vivo* oligoTRAFTAC activity. Overall, our data demonstrate oligoTRAFTACs as a generalizable platform towards difficult-to-drug TFs and their degradability via the proteasomal pathway.

## INTRODUCTION

Protein degradation is an integral component of cellular homeostasis which supports the healthy environment within the cell. The ubiquitin-proteasomal pathway is one of the key mechanisms by which unwanted or defective proteins are degraded (*1, 2*). Targeted protein degradation by PROteolysis TArgeting Chimeras, or PROTACs, has stimulated the development of novel strategies to induce protein degradation, for both discovery biology and therapeutics (*3*). PROTACs are small molecule-based heterobifunctional molecules that recruit an E3 ligase complex to a protein of interest (POI) (*4*). By doing so, PROTACs induce the ubiquitination and degradation of the POI via the proteasome. Although PROTACs have the potential to induce the degradation of numerous proteins, the identification of small molecule recruiting ligands for several classes of proteins is still challenging and time-consuming (*5*). Therefore, the development of alternative, proximity-inducing strategies would help to target disease-causing proteins, such as transcription factors.

Transcription factors (TFs) control gene expression by binding to specific DNA elements within a given gene promoter or distal enhancer regions (*6, 7*). Many diseases such as neurological disorders, autoimmunity, developmental syndromes, and many cancers result from abnormalities in the TF-controlled gene-regulatory circuitry within the diseased cell (*8-10*). For instance, the TF c-Myc, the most frequently amplified oncogene, has been extensively studied and established as a direct mediator of tumorigenesis in numerous cancers (*11*). Although indirect approaches such as bromodomain protein inhibitors and PROTACs have been explored to control c-Myc levels, direct inhibition or degradation of c-Myc has been an as yet unrealized goal (*12, 13*). In contrast to c-Myc, T-box transcription factor (“brachyury”) is expressed only in a restricted set of cancer types and minimally expressed in normal cells (*14*). Brachyury is a master developmental TF that plays a pivotal role during early embryonic development in vertebrates (*15*). Brachyury expression is limited only to embryonic developmental stages and, in general, expression levels are highly downregulated in adult tissues (*16*). In addition to its regulatory functions during development, brachyury expression in remnant notochord cells in adults has been shown to be a key oncogenic driver in the rare bone cancer, chordoma (*17*). Chromosomal aberrations such as chromosome 6 gains and partial polysomy have been identified as potential molecular mechanisms in brachyury-dependent chordoma tumors (*18-21*). Other TFs such as NF-kB, STAT3/5, the androgen receptor (AR) and the estrogen receptor (ER) are other known oncogenic drivers that also rewire transcriptional circuitry in various cancer types (*22, 23*). Although hundreds of TFs have been identified as key players in human diseases, the lack of direct therapeutic strategies is a bottleneck for accessing these difficult-to-drug targets.

Although several TFs, such as STAT3, AR and ER have been successfully degraded by PROTACs, a plethora of disease-relevant yet traditionally “undruggable” TFs remain unaddressed (*24-26*). Recently, we developed a generalizable TF degradation platform by taking advantage of the DNA binding ability of TFs (*27, 28*). In this approach, we generated a chimeric DNA:CRISPR RNA molecule -- a TRAnscription Factor TArgeting Chimera (TRAFTAC) -- that binds a dCas9-HaloTag7 fusion adaptor, which in turn recruits an E3 ligase into proximity with the transcription factor of interest (TOI). Using TRAFTACs, we demonstrated for the first time that TF response elements could be used in degrader design to recruit and induce the degradation of TOI, while highlighting their generalizability as well. While that strategy could be readily used as a chemical biology tool to study the biology of understudied TFs, therapeutic application of the technology is limited.

In this current report, we describe the development of second-generation TRAFTACs – “oligoTRAFTACs” that are based solely on an oligonucleotide sequence and E3 ligase-recruiting small molecule ligand to recruit TOI and the cellular ubiquitination machinery, respectively. A short oligonucleotide specific to a TOI was synthesized with a terminal alkyne either at the 3’ or 5’ end. Copper-catalyzed alkyne-azide cycloaddition (CuAAC) click reaction between the alkyne-oligonucleotide and an azide-containing VHL ligand yielded chimeric oligoTRAFTACs that enabled c-Myc and brachyury degradation in HeLa and chordoma cell lines, respectively. Furthermore, based on zebrafish experiments, we demonstrate the feasibility of oligoTRAFTACs application to *in vivo* settings without the requirement for genetic modification (i.e., ectopic expression) in the system.

## Results and Discussion

### Degradation of c-Myc by oligoTRAFTACs

Myc transcription factors are dysregulated in a range of cancers. Even though c-Myc has been targeted by indirect approaches, development of direct-targeting methods is hindered by the paucity of ligandable pockets and their highly disordered structure (*29*). Since c-Myc binds to specific DNA sequences with a conserved E-box sequence (CACGTG), we sought to incorporate a c-Myc binding consensus DNA-sequence into the oligoTRAFTAC (OT) design (*30*). We adapted a Myc binding consensus sequence (5’ **CACGTGGTTGCCACGTG 3’)** taken from one of its target gene promoters (*31*). Additionally, we included a flanking sequence at both 3’ and 5’ end of the c-Myc-targeting oligonucleotide sequence to facilitate successful double strand formation of the recognition sequence while providing a flexibility for oligoTRAFTACs. First, we synthesized a c-Myc-targeting oligonucleotide sequence with an orthogonal alkyne handle on either side as a reactive moiety to append an azide-containing VHL ligand (Figure S1 A). Copper-catalyzed alkyne-azide cycloaddition (CuAAC) click reaction was performed to synthesize two c-Myc targeting oligoTRAFTACs (OT7: VHL ligand at the 5’ end and OT10: VHL ligand at the 3’ end of the oligonucleotide) (Figure S1 B, C). After 18 h at room temperature, the crude mixture was purified and analyzed by HPLC to verify reaction completion (Figure S2 A). Furthermore, the reaction was also monitored by electrophoretic mobility shift assay (EMSA) (Figure S2B). The single stranded oligoTRAFTACs were then purified by HPLC, dried and reconstituted in water. To generate double stranded oligoTRAFTACs, we then performed an annealing reaction in the presence of the reverse complementary oligonucleotide by heating to 95 ^0^C and slowly cooling to room temperature over a 1.5 h period. In addition, we also synthesized a biotinylated version of the same c-Myc-targeting oligonucleotide to generate a double stranded oligonucleotide via the same annealing reaction conditions.

First, we performed a biotin-pulldown experiment with the c-Myc-targeting biotin-oligonucleotide and scrambled control to confirm that the selected oligonucleotide sequence binds c-Myc. After incubation of biotin probes with HeLa cell lysate and their capture using streptavidin agarose, the eluates were probed with c-Myc antibody (Figure 2A). The data indicated successful c-Myc engagement with the biotin-oligonucleotide probe compared to its scrambled version, thereby predicting c-Myc recruitment by the oligoTRAFTAC. We next set out to test c-Myc degradation upon transfection of increasing concentration of OT7 into HeLa cells. After 20 h post transfection, western blot analysis of the cell lysates indicated that OT7 can induce significant c-Myc degradation at 50 nM (Figure 2B). A similar pattern was observed in HEK293T cells in response to OT7 transfection (Figure S2C). To test the kinetics of oligoTRAFTAC-mediated c-Myc degradation we performed a post-transfection washout experiment. After 6 h or 12 h post-transfection, cell culture medium was replaced with fresh medium and incubated for a total of 20 h. When cells were transfected for 6 h prior to washout, we observed very low c-Myc degradation levels relative to the 12 h transfection (Figure S2D). Furthermore, the degradation was maintained for 24 h (total of 36 h) post-washout indicating a possible catalytic behavior of oligoTRAFTACs (Figure S2E).

**Figure 1.**
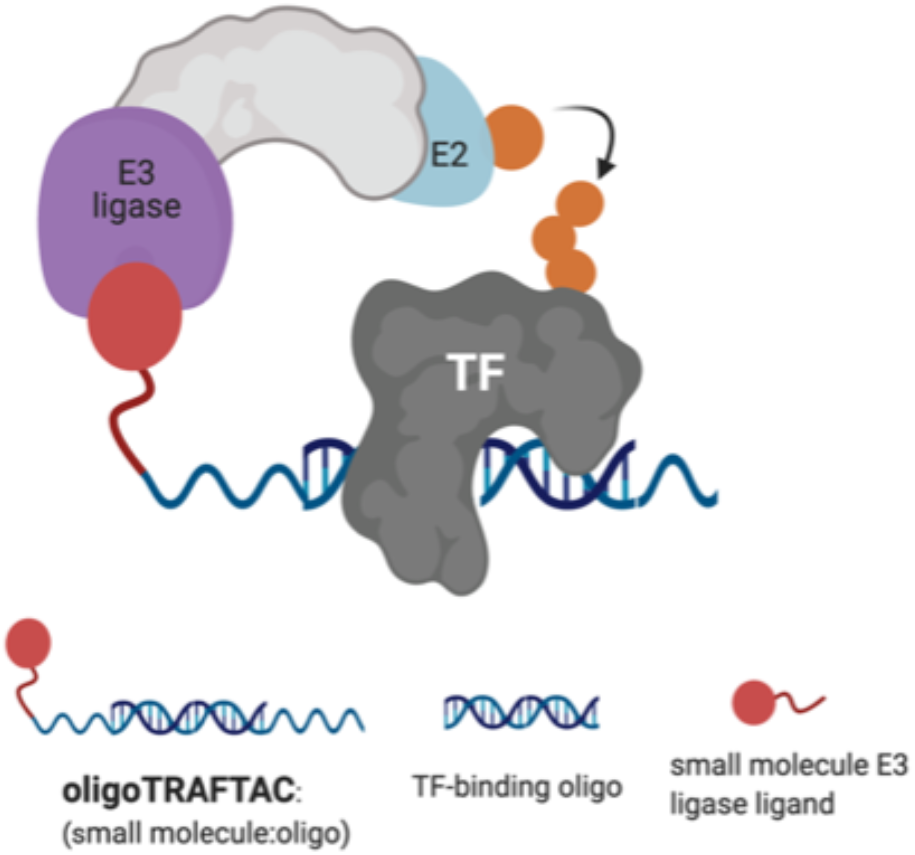
Schematic representation of oligoTRAFTAC-mediated transcription factor (TF) and E3 ligase recruitment, and proximity-dependent TF ubiquitination.

**Figure 2.**
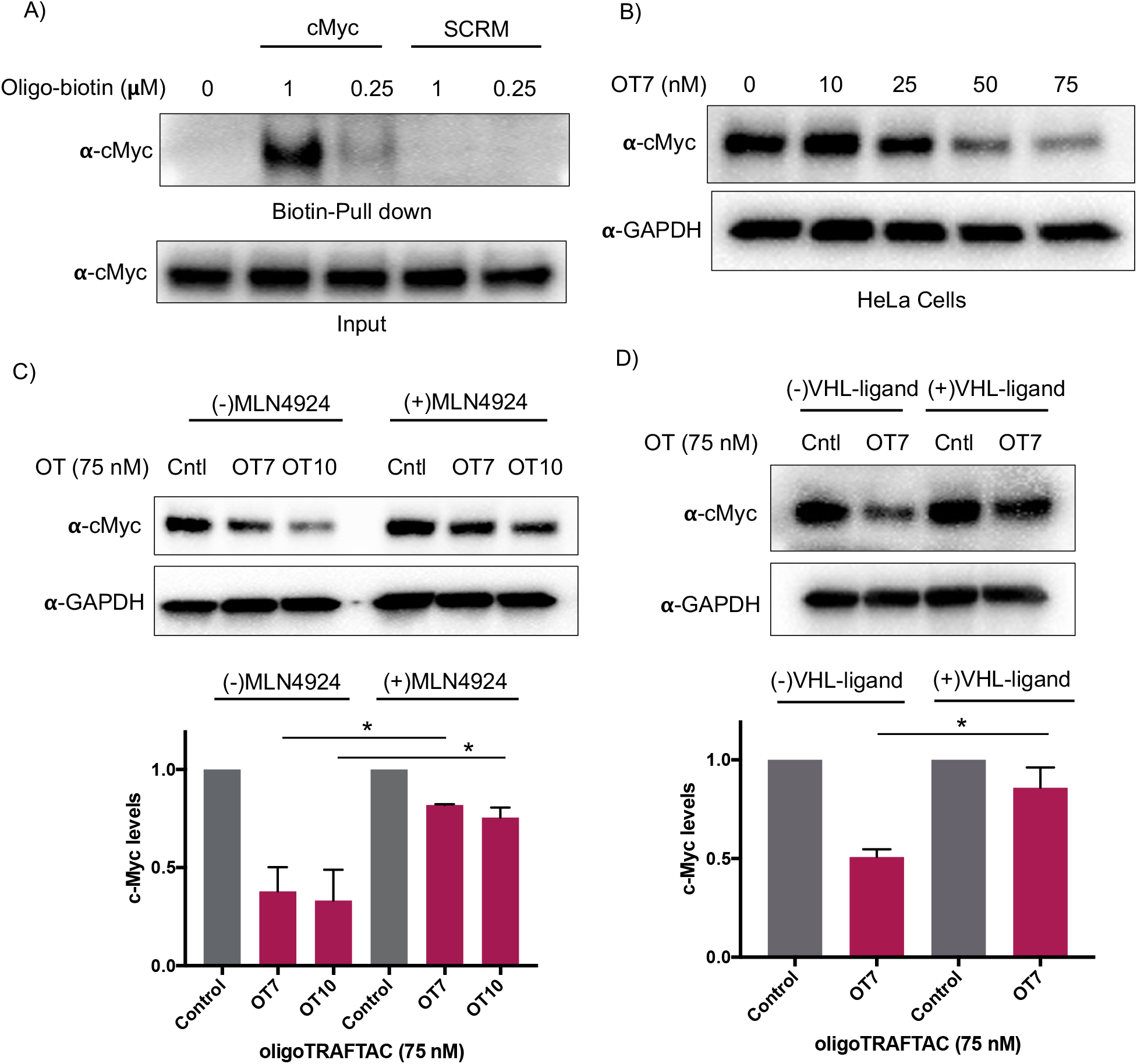
OligoTRAFTAC induces c-Myc degradation. A) The oligonucleotide selected for the c-Myc oligoTRAFTAC engages c-Myc. HeLa cell lysates were incubated with biotinylated oligonucleotide, or its scrambled sequence, followed by capture with streptavidin agarose and probing for c-Myc. B) Dose response of OT7-mediated c-Myc knockdown in HeLa cells. C) oligoTRAFTAC-induced c-Myc degradation occurred via the proteasomal pathway. HEK293T cells were treated with c-Myc-targeting oligoTRAFTAC with and without the neddylation inhibitor, MLN-4924 (1 µ M), and then analyzed for c-Myc levels. D) HEK293 cells were preincubated with and without 10 µM VHL ligand followed by OT7 transfection and analyzed for c-Myc levels.

After confirming that c-Myc is susceptible to oligoTRAFTAC-mediated degradation, next we tested whether the observed degradation occurs through the proteasomal pathway. To address this, we evaluated whether neddylation inhibition, which disrupts cullin RING E3 ligase function, affects the activity of the c-Myc-targeting oligoTRAFTACs. When cells were pre-incubated with MLN-4924, c-Myc degradation was significantly diminished, demonstrating that OT7 and OT10-mediated c-Myc degradation is neddylation-dependent and occurs via the proteasomal pathway (Figure 2C). Next, we performed a VHL ligand competition experiment to confirm that oligoTRAFTAC-mediated c-Myc degradation is VHL-dependent. Cells were preincubated in the presence or absence of excess VHL ligand for 1.5 h before transfecting oligoTRAFTACs for 20 h. As anticipated, we observed c-Myc levels in the VHL ligand-incubated cells were maintained relative to the cells that were not incubated with VHL ligand (Figure 2D). Overall, the data indicated that OT7 and OT10 induces c-Myc degradation via the proteasomal pathway. To test the effect of c-Myc degradation on cell proliferation, we assessed the cell viability in response to the transfection of OT7 and OT12. As the data indicate, the induction of OT7-mediated c-Myc degradation inhibited cell proliferation compared to scrambled control-transfected cells (Figure S2F).

### Brachyury targeting oligoTRAFTACs mediated brachyury degradation

T-box transcription factor (brachyury) is a DNA-interacting TF that regulates gene expression during early embryonic development in vertebrates. Brachyury is crucial for early mesoderm formation and plays a key role in notochord development where, as a homodimer, it recognizes and binds to a consensus palindromic DNA sequence (*32-34*). To investigate brachyury degradation by oligoTRAFTACs, we adapted a brachyury-binding DNA sequence (AATTTCACACCTAGGTGTGAAATT) as the brachyury-recruiting element in our new oligoTRAFTACs design (*35*). We synthesized oligonucleotides with flanking bases and terminal alkyne moieties at either end (3’ and 5’) of the oligonucleotide (Figure S1A, right panel).

To generate brachyury-targeting oligoTRAFTACs, we synthesized two azido-VHL ligands with a long (5 PEG units) and a short (2 PEG units) linker (Figure S1C, S3A). We tested whether the oligonucleotide used in oligoTRAFTAC design could recruit brachyury by performing a streptavidin pull-down experiment using a biotin-oligonucleotide. After incubating the brachyury targeting biotin-oligonucleotide or its scrambled version for 1.5 h with the lysate of HEK293T cells that stably express a brachyury-GFP fusion, streptavidin beads were added to cell lysates and incubated for another 16 h at 4 °C to capture the binary complex. After several washings, eluted fractions were analyzed by western blotting. Biotin pulldown data indicated that the oligonucleotide used in brachyury-targeting oligoTRAFTAC design is engaged with its target protein (Figure 3A). In contrast, scrambled oligonucleotide failed to bind brachyury-GFP, indicating sequence-specific brachyury recruitment.

**Figure 3.**
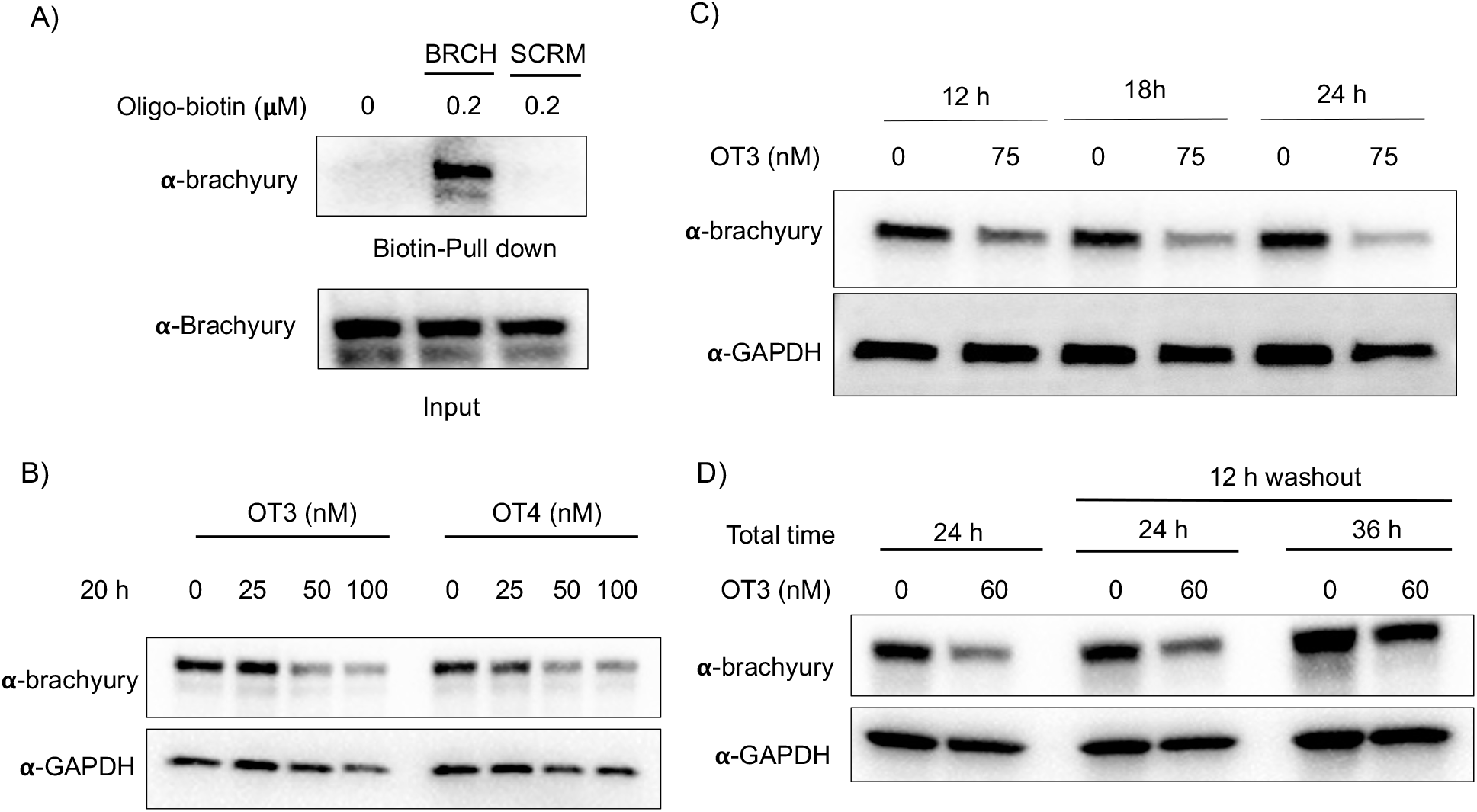
Brachyury-GFP degradation by oligoTRAFTACs. A) Brachyury-targeting oligonucleotide used in the oligoTRAFTAC design engaged with brachyury-GFP. Brachyury targeting biotinylated oligonucleotide (BRCH) or its scrambled oligonucleotide (SCRM) incubated with cell lysate and captured by streptavidin agarose beads. B) Two oligoTRAFTACs with 3’ VHL ligand modifications, OT3 (5 PEG unit linker) and OT4 (2 PEG unit linker) were transfected into HEK293T cells and brachyury-GFP levels were analyzed in lysates prepared after 20 h. C) OT3 induced brachyury-GFP degradation as early as 12 h in HEK293T cells. D) Washout experiment after 12 h of OT3 transfection. Cells were incubated continuously for 24 h in transfection medium or OT3 was aspirated after 12 h of transfection and fresh medium added to cells. Washout cells incubated for another 12 h and 24 h prior to harvesting.

To test whether a brachyury-targeting oligoTRAFTAC induces degradation, brachyury-oligoTRAFTACs were transfected into brachyury-GFP-expressing HEK293T cells for 24 h, after which cells were collected and lysed, followed by western blotting. OligoTRAFTACs with 5’ VHL-ligand and a longer linker (OT1) did not induce significant degradation of brachyury-GFP (Figure S3B), while its counterpart the shorter linker (OT2) successfully induced target degradation. However, oligoTRAFTACs with 3’ VHL-ligand (OT3 and OT4) both showed comparable, better degradation profiles relative to OT1 and OT2 (Figure S3B last two panels). Next, different concentrations of OT3 and OT4 were transfected into HEK293T cells and lysed after 20 h. Data indicated that both OT3 and OT4 could induce significant brachyury-GFP degradation at 50 nM concentration within 20 h of transfection (Figure 3B). Furthermore, time course experiment suggested that OT3 can induce significant brachyury-GFP degradation at 12 h that is maintained up to 36 h post-transfection (Figure 3C and Figure S4A). Next, we performed a washout experiment to determine the minimum incubation time to induce a noticeable brachyury degradation by OT3. To address this question, we transfected 75 nM of OT3 into HEK293T cells expressing brachyury-GFP followed by replacement of the transfection medium after 6 h and 12 h with fresh cell culture medium (Figure S4B). Degradation data indicated that OT3 incubated for at least 12 h induces significant brachyury degradation comparable to continuous 24 h incubation, whereas OT3 incubation for 6 h did not induce significant brachyury degradation (Figure 3D, center panel). Furthermore, brachyury degradation was prominent even after the cells incubated in fresh medium for longer time (36 h), which is consistent with oligoTRAFTAC OT7 targeting c-Myc-targeting (Figure 3D, last panel). However, brachyury levels at 36 h had increased relative to the 24 h period, indicating a progressive loss of brachyury degradability by OT3. This could be partially explained by the reduced stability of oligoTRAFTACs due to the oligonucleotide sensitivity towards intracellular nucleases, increased deubiquitinase (DUB) activity, or increased c-Myc resynthesis.

To test whether the oligoTRAFTAC-mediated brachyury recruitment and degradation is oligonucleotide-dependent, we synthesized and evaluated a scrambled-oligoTRAFTACs (OT5 and OT6) for brachyury degradation in cells. Consistently, OT3 induced a robust brachyury-GFP degradation, whereas neither OT5 nor OT6 induced significant brachyury degradation (Figure 4A, Figure S5). We next performed a VHL ligand competition experiment to confirm brachyury degradation is dependent on VHL. In this experiment, cells were pretreated with 100-fold excess VHL ligand prior to OT3 treatment. As anticipated, VHL competition rescued oligoTRAFTAC-mediated brachyury degradation (Figure 4B), confirming that observed brachyury degradation is a result of VHL E3 ligase recruitment by OT3. Furthermore, we evaluated OT3-mediated brachyury degradation in the presence of a neddylation inhibitor to further confirm that the intended mechanism is via the proteasomal pathway. Similar to the VHL competition experiment, the neddylation inhibitor, MLN-4924 was pre-incubated with cells at 1 µM concentration. After 1.5 h, OT3 was transfected into the cell for 20 h following which the cell lysates were analyzed as indicated in Figure 4C. The data indicated that OT3 could not induce brachyury degradation in the presence of MLN-4924, showing that neddylation is crucial for the mechanism of action (Figure 4C). Furthermore, analysis of GFP-fluorescence confirmed brachyury degradation only in the absence of MLN-4924 (Figure 4D). Overall, these data support the fact that OT3-induced brachyury degradation is mediated through the proteasomal pathway and is dependent on the oligonucleotide sequence of OT3.

**Figure 4.**
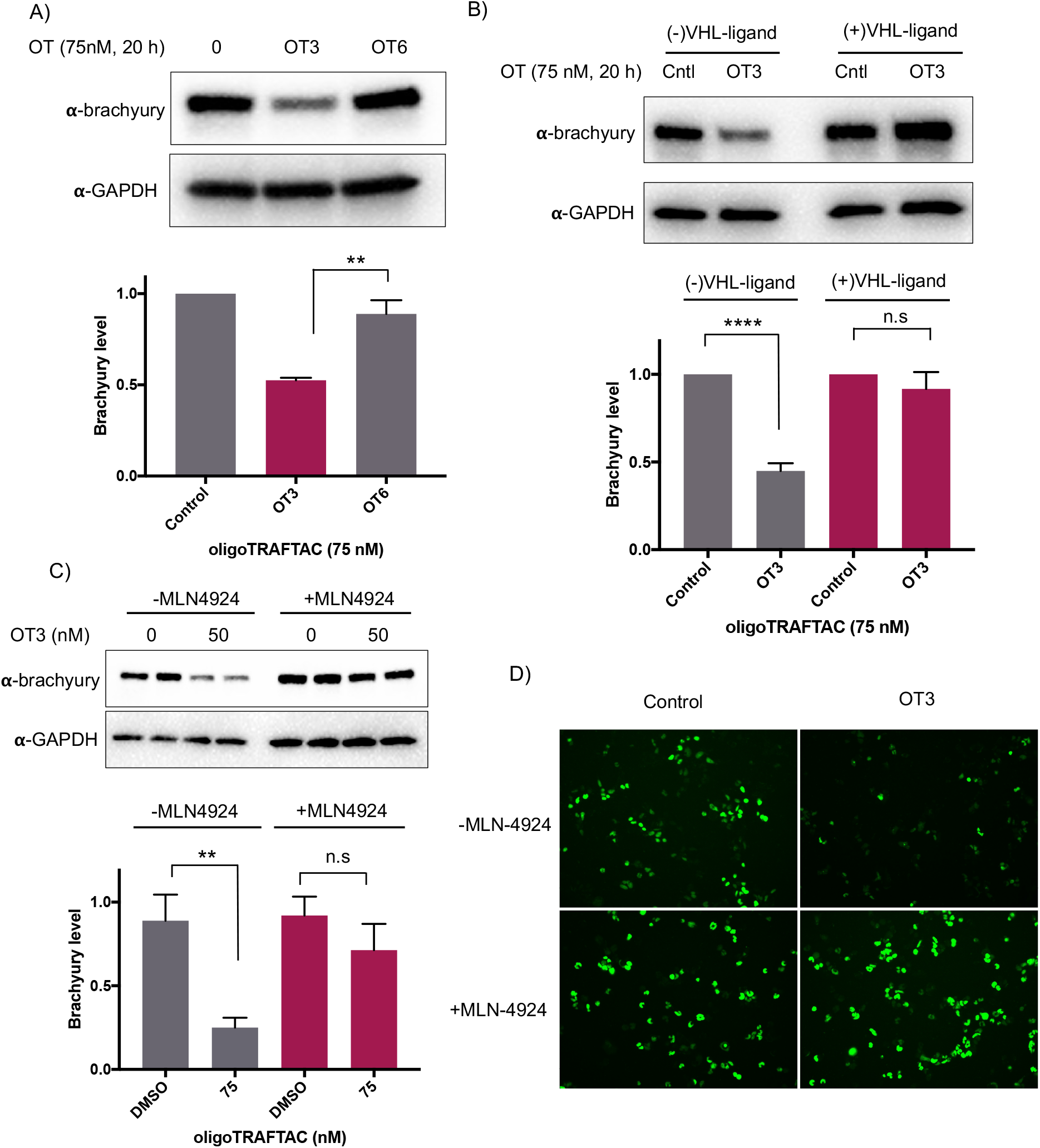
OligoTRAFTACs induce brachyury-GFP degradation via the proteasomal pathway. **A)** HEK293T cells expressing brachyury-GFP were transfected with 75 nM each of OT3 and its scrambled OT6, and lysates were probed for brachyury levels. B) OT3 induced brachyury degradation is VHL-dependent. HEK293T cells were preincubated with and without 10 µM of VHL ligand for 1.5 h prior to OT3 transfection. After 20 h of transfection, cells lysates were prepared and analyzed for brachyury degradation. C) OT3 induces brachyury degradation via the proteasomal pathway: neddylation inhibitor MLN-4924 was preincubated with cells prior to OT3 transfection. After 20 h of transfection of OT3, cells were harvested and analyzed for brachyury levels. D) Brachyury-GFP downregulation was monitored by GFP fluorescence in cells in the presence or absence of MLN-4924.

### OligoTRAFTACs induce brachyury degradation in chordoma cells

Increasing OT3 concentrations were transfected into UM-Chor1 cells followed by western blotting to determine brachyury degradation. Consistent with the brachyury-GFP degradation in HEK293T cells, 60 nM of OT3 induces ∼70% brachyury degradation in UM-Chor1 cells 24 h post-transfection (Figure S5B). Although OT3 induced comparable levels of brachyury degradation in both cell lines, phosphodiester linkages within OT3 are susceptible to cleavage by both extra- and intracellular nucleases. Therefore, we next synthesized oligoTRAFTACs with a phosphorothioate (PS) backbone (*36*). The addition of extra non-bridging sulfur atoms in the inter-nucleotide phosphate group has been shown to have both increased stability against nucleases and improved cell permeability (*36*). A brachyury-targeting oligoTRAFTAC, OT17, was synthesized by incorporating PS bonds throughout the oligo sequence. Increasing OT17 concentrations were transfected into chordoma cells (UM-Chor1) for 24 h following which the cells were lysed and probed for brachyury levels -- western blotting showed that OT17 induced significant degradation of endogenous brachyury even at 15 nM (Figure 5A). We noticed a similar degradation pattern in another chordoma cell line, JHC-7 (Figure 5B), although OT17 was slightly less potent in them, requiring 30 nM to induce brachyury knockdown comparable to UM-Chor1 cells. Time course experiment showed that OT17 induced a modest degradation at 8 h, which increased to ∼ 60% knockdown 16 h post-transfection (Figure 5C) although washout experiment indicated that incubation of transfection complex for 12 h is sufficient to induce significant brachyury degradation (Figure 5D). Furthermore, this same experiment displayed persistent degradation for 24 hours post washout (total of 36 h of post-transfection), suggesting an increased stability and possible catalytic mechanism of OT17. We also synthesized and tested a scrambled oligoTRAFTAC with a PS backbone throughout the oligonucleotide sequence (OT20). Consistently, side-by-side comparison of brachyury levels UM-Chor1 in cells transfected with OT17 and OT20 indicated sequence-specific brachyury degradation (Figure 5E). To monitor oligoTRAFTAC-mediated brachyury ubiquitination resulting from induced proximity to VHL, we next transfected HA-ubiquitin into HEK293 cells overexpressing brachyury-GFP. After cells pre-incubated with proteasome inhibitor epoxomicin (500 nM), OT3 and OT17 were transfected and incubated for 12 hours. Subsequent immunoprecipitation data indicated that OT3 and OT17 could induce ubiquitination of brachyury-GFP, confirming that degradation by oligoTRAFTAC follows a ubiquitination event mediated by recruited VHL (Figure 5F).

**Figure 5.**
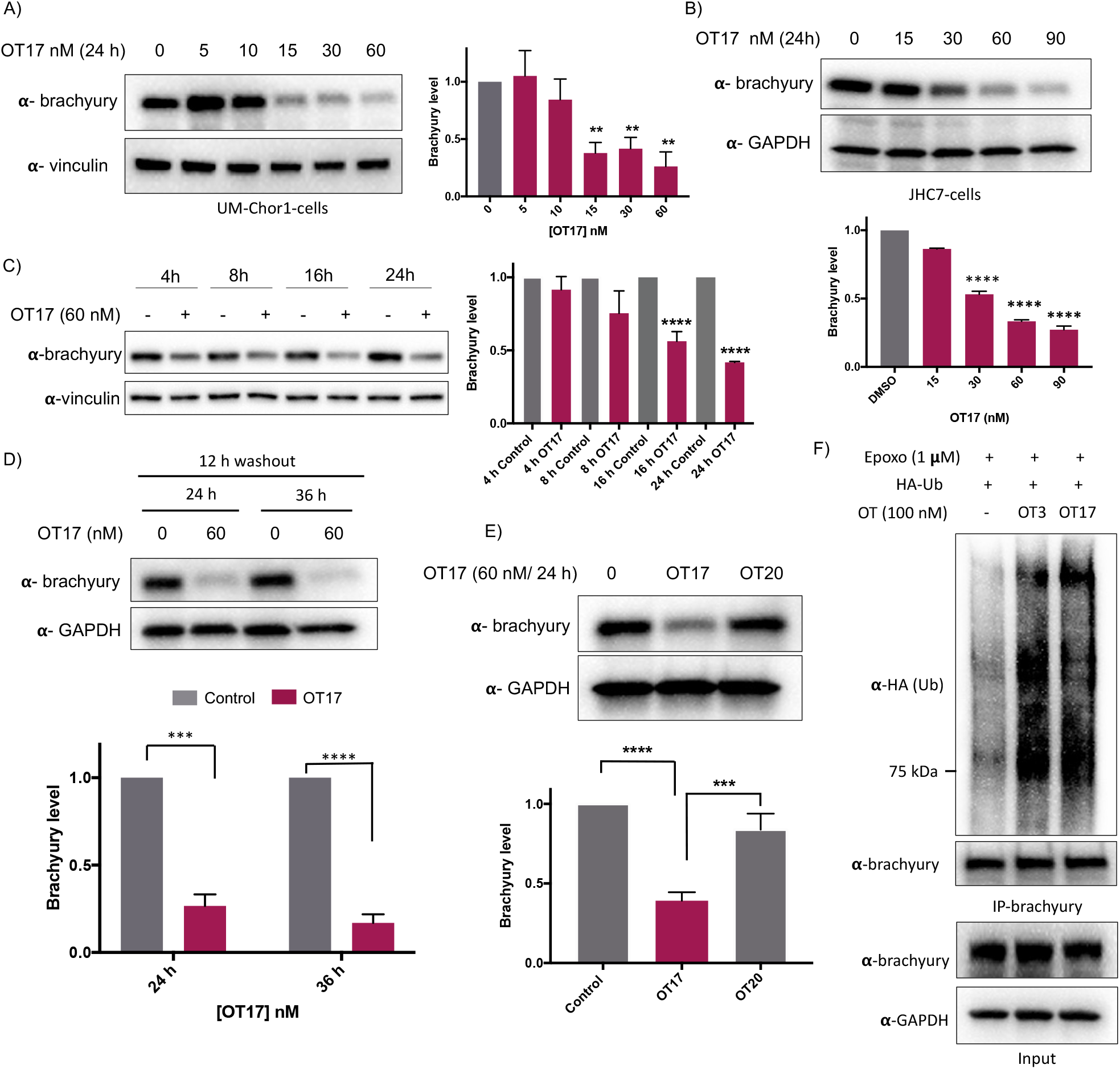
Endogenous brachyury degradation by oligoTRAFTACs constructed with phosphorothioate backbone. A) Increasing concentrations of OT17 were transfected into UM-Chor1 cells and harvested after 24 h, subjected to lysis and analyzed for brachyury downregulation. Brachyury levels were normalized to loading control and presented as a bar graph. B) JHC-7 cells were transfected with OT17 and probe for brachyury levels as explained in (A). C) UM-Chor1 cells were transfected with 60 nM of OT17 and harvested at subsequent different time points as indicated. D) Washout experiment: transfection medium was removed after 12 h of OT17 transfection and UM-Chor1 cells were incubated for another 12 h or 24 h in fresh complete cell culture medium. E) OT17-mediated brachyury degradation is oligonucleotide sequence dependent. UM-Chor1 cells were transfected with OT17 and scrambled OT20, cells were lysed and analyzed as shown. F) OT3- and OT17-induced brachyury ubiquitination. HEK293T cells that overexpress brachyury-GFP were transfected with HA-ubiquitin, followed by the second transfection with OT3 or OT17. After 12 h, cell lysates were subjected to immunoprecipitation using brachyury antibody, and the eluates blotted for the indicated proteins.

### *In vivo* activity of oligoTRAFTACs: OT17-mediated developmental defects in zebrafish tail formation

After confirming that brachyury-targeting oligoTRAFTACs induce efficient endogenous brachyury degradation in cells, we next evaluated their activity in animals using zebrafish as a model organism. Although brachyury overexpression in adult tissue is one of the key factors that leads to tumorigenesis in chordoma, brachyury is widely known for its essential biological activity in vertebrate notochord formation at early stages of embryonic development. To test for oligoTRAFTAC *in vivo* activity, we examined tail deformation in OT3-injected zebrafish embryos relative to mock-injected embryos (Figure 6A). Interestingly, although OT3 could induce brachyury degradation in cells, OT3 did not induce tail deformation in zebrafish, possibly due to the sensitivity of its phosphodiester backbone to nucleases. However, microinjection of OT17, in which PS linkages provide stability against both exonucleases and endonucleases, induced tail deformation in ∼70% of injected embryos (Figure 6B, C) whereas <5% mock and scrambled OT20 injected embryos had defective tails. This result illustrates how the observed brachyury loss of function phenotype is sequence-dependent, and demonstrates *in vivo* oligoTRAFTAC activity in zebrafish.

**Figure 6.**
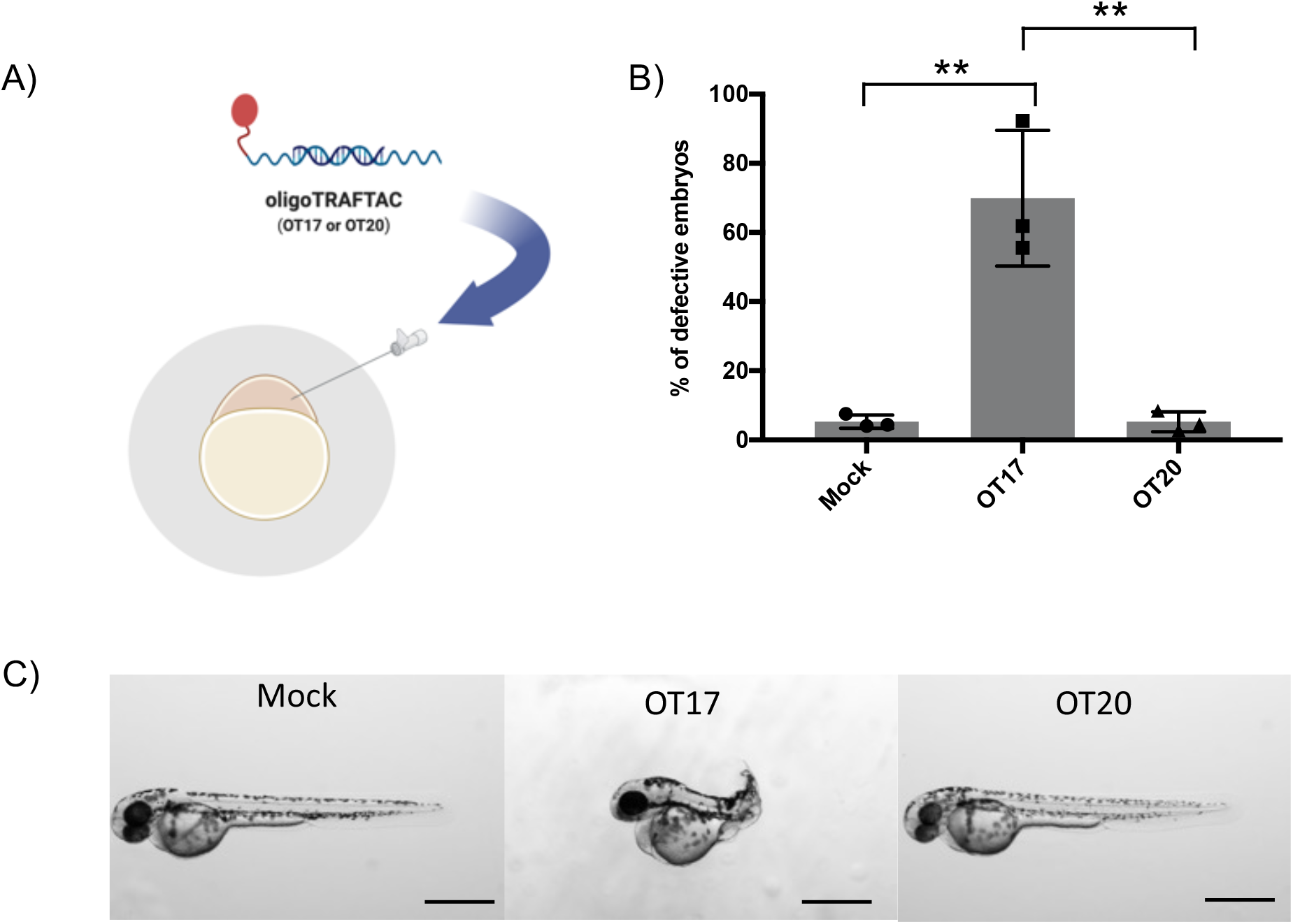
Microinjection of brachyury-targeting oligoTRAFTAC into zebrafish embryos: demonstration of *in vivo* activity. A) Schematic representation of OT17 and OT20 microinjection into zebrafish embryos. B) Mock, OT17 and OT20 (180 picoliters from 25 µM of oligoTRAFTACs, or mock equivalent) were microinjected into embryos (number of embryos in each group for three independent experiments; mock-47, 50, 43; OT17-49, 52, 61; OT20-75, 74, 45).After 48 h, the number of defective tails in each group was recorded and presented as percentage in a bar graph. C) Images of representative zebrafish from the cognate treatment groups. Pictures were captured after 48 h post microinjection of mock, OT17 and OT20. Scale bar 500 µm.

## Discussion

The majority of transcription factors, key mediators of gene expression, are considered undruggable. In this study, we developed a method for oligonucleotide-dependent TF recruitment and degradation by the proteasomal pathway. We have coupled TF-binding short DNA sequences from target gene promoters with VHL ligands to create bifunctional molecules for the targeted degradation of those same TFs. Since the binding sequences already have been identified for many TFs, and since synthetic routes for oligonucleotide synthesis are well established and economical, these oligonucleotide sequences can be rapidly employed in oligoTRAFTAC design to use as a versatile tool for both basic discovery biology and therapy development (*37*). We previously developed a strategy (first generation TRAFTACs) based on using an oligonucleotide and E3 ligase recruiting intermediate protein, dCas9-Halotag7, by exploiting its ability to recruit both TOI and E3 ligase in the presence of TRAFTAC and a concurrently administered HaloPROTAC (*27*). Following the first generation TRAFTAC study, a couple of other studies were published describing the use of oligonucleotides for TF degradation (*38, 39*). Although first generation TRAFTACs are widely applicable as chemical biology tools for discovery biology, therapeutic applicability needed further improvements. In this current study, we discuss the development of second generation TRAFTACs by replacing the intermediatory E3 ligase recruiting dCas9-Halotag7 in the first generation of TRAFTACs with a VHL-recruiting small molecule ligand.

Second generation TRAFTACs were synthesized by directly attaching a VHL ligand to c-Myc or brachyury-binding oligonucleotide sequences via click chemistry. OligoTRAFTACs targeting c-Myc TF displayed a robust degradation. Addition of VHL ligand to either side of the oligonucleotide did not significantly alter its ability to induce c-Myc degradation. This can be partially attributed to the flexibility of oligoTRAFTACs provided by the extra flanking nucleotides between the VHL ligand and TF-recruiting oligonucleotide. Other oligoTRAFTACs used in this study induced the degradation of ectopically expressed brachyury-GFP and endogenous brachyury via the proteasomal pathway. Although washout experiments indicated that oligoTRAFTACs are not as kinetically efficient as conventional PROTACs, the transient binding nature of oligoTRAFTACs to both TF and VHL proteins suggests the probability of a similar catalytic degradation mechanism. Moreover, to our knowledge the current study provides the first evidence of the degradability of untagged, endogenous brachyury in multiple chordoma cell lines. In the first attempt to test oligoTRAFTAC *in vivo* activity, OT3 (containing a phosphodiester backbone) could not induce the intended defective tail phenotype in zebrafish. It is noteworthy that first generation TRAFTACs, where oligonucleotides are partly protected by the ribonuclearcomplex, induced defective tails in zebrafish.

Therefore, a potential reason for the failure of OT3 to induce the intended phenotype within the embryos may be their high sensitivity and exposure of oligoTRAFTACs to embryonic nucleases present at the early stages of development. Therefore, phosphodiester oligoTRAFTACs might not endure *in vivo* at the intracellular concentrations needed to achieve a significant brachyury degradation and the intended phenotype. Previous studies suggested that synthesis of oligonucleotide-based entities with a phosphorothioate backbone largely benefitted from nuclease resistance. Therefore, the microinjection of phosphorothioate backbone-containing OT17 resulted in a significantly high occurrence rate of the defective tail phenotype compared to the mock or scrambled oligoTRAFTAC (OT20), confirming the increased stability and resistance of OT17 to nucleases. Overall, oligoTRAFTACs are programmable heterobifunctional molecules comprised of a TF-binding oligonucleotide sequence and a VHL binding ligand, which induce TF degradation in cells and displayed robust *in vivo* activity. Due to the simple and modular nature of their structure, oligoTRAFTACs can be rapidly designed for many non-ligandable DNA-binding TFs both for use as a chemical biology tool as well as a potential therapeutic strategy.

## ACKNOWLEDGEMENT

We thank Dr. John Hines and Dr. Dhanusha A. Nalawansha for the critical reading of the manuscript, insightful comments, and critiques. C.M.C. is funded by the NIH (R35CA197589) and is supported by an American Cancer Research Professorship. Funding provided by NIH grants R01GM129149 to S.A.H and 1F32GM127502 to M.A.G.

## Supporting Information

## Materials and Methods

### Materials and reagents

All cell culture media and fetal bovine serum (DMEM, IMDM, RPMI, DMEM/F-12) were purchased from Gibco unless otherwise specified. Primary antibodies for GAPDH (2118S), Vinculin (13901S), brachyury (81694S), HA-tag (3724S), protein A magnetic beads (73778S) and streptavidin magnetic beads (5947S) were purchased from Cell Signaling Technologies. Primary antibody for c-Myc (sc-40) was purchased from Santa Cruz. Secondary rabbit (NA934) and mouse (NA931) antibodies were purchased from GE Health Care. RNAiMAX (13778-150) transfecting reagent was purchased from Thermo Fisher Scientific. MLN4924 (S7109) was purchased from Selleckchem. Copper (II) sulfate pentahydrate (209198) and L-Ascorbic acid (A4403) were purchased from Millipore Sigma, and THPTA purchased from Lumiprobe (H4050). All the oligonucleotide modifiers were purchased from Glen Research, 3’ Alkyne (20-2992-41), 5’ Alkyne (10-1992-90) and biotin (10-5950-90). All the oligonucleotides were custom synthesized by Yale Keck oligo synthesis facility.

### General biology methods

#### Cell culture

Human embryonic kidney cells HEK293T cells and HeLa cells were grown in Dulbecco’s Modified Eagles Medium (DMEM) containing 10% heat inactivated fetal bovine serum (FBS), streptomycin (5 µg/mL) and 5 U/mL penicillin 95 U/mL). All cell lines were maintained and cell culture experiments were carried out in humidified incubators at 37 degrees and 5% CO_2_ supplementation.

### OligoTRAFTAC transfection

One day prior to oligoTRAFTACs transfection, cells (HEK293T: 0.7×10^6^/well, HeLa: 0.2 ×10^6^/well, UM-Chor1: 0.2×10^6^/well and JHC7: 0.4×10^6^/well) were propagated into 6-well plates containing appropriate complete growth medium. Prior to transfection, complete medium was replaced with 1.75 mL of transfection medium (2%FBS, no Penstrep). Chimeric oligoTRAFTAC transfection was performed using RNAi-Max reagent according to the protocols provided by the manufacture. All transfections were carried out in 6-well plates with 2 mL of media and concentrations of oligoTRAFTACs were calculated according to this volume (2 mL). Briefly, for 50 nM concentration, 4 µl from a 25 µ M oligo TRAFTAC stock was added to a tube containing 125 µl of OPTIM-MEM and 12.5 µl of RNAi-Max reagent was added to a separate tube containing 125 µl of OPTIM-MEM (added ∼4µg of oligoTRAFTAC to 2 mL cell culture medium). Two tubes were incubated for 5 minutes at room temperature and oligoTRAFTAC containing OPTI-MEM was then slowly added to the second tube with RNAi-MAX. The solution in the tube was mixed well by pipetting up and down several times. After incubating for 10 minutes at room temperature, 250 µl of oligoTRAFTAC:RNAi-MAX complex was added drop wise onto cells containing the transfection medium. Transfection medium was mixed well before transferring the 6-well plate into the incubator. After appropriate time, cells were either harvested or transfection medium containing oligoTRAFTAC:RNAi-MAX complex was replaced with fresh medium and incubated for desired time point prior to harvesting. For MLN-4924 and VHL ligand competition assay, these molecules were pre-incubated in the transfection medium (1.75 mL) for 1 h prior to transfection of oligoTRAFTACs. Cell lysates were prepared by incubating cells in RIPA lysis buffer (25 mM Tris pH 7.6, 150 mM NaCl, 1% NP40, 1% deoxycholate, 0.1% SDS, 1X protease inhibitor cocktail from Roche and 1 mM of PMSF) on ice for 30 minutes and cell lysate was clarified by centrifugation at high speed (15 000 rpm) for 20 minutes. Clear supernatant was collected for further experiments. For cell proliferation/viability assays, cells were split (HeLa: 0.1 ×10^5^/well) into white, clear bottom 96-well plates. Transfection was carried out as described above, except volumes, i.e., 175 µl of transfection medium per one well, 0.5 µl of RNAi-MAX/well and 25 µl of OPTIMEM per well.

### Click rection

Alkyne-modified oligonucleotides were dissolved in ultra-pure water at 500 µ M concentrations and azide-modified VHL Ligands were dissolved in DMSO at 10 mM. Right before the click reaction, fresh stock solutions of Cu (II) sulfate pentahydrate (50 mM in water), Tris(3-hydroxypropyltriazolylmethyl)amine (THPTA: 100 mM in DMSO) and sodium ascorbate (100 mM in water) were made. The click reaction was carried out in 50% DMSO solution. First, alkyne modified oligonucleotide (250µl) and azide-modified VHL Ligand (1:5 molar ratio) mixed in tube 1. Then, Cu (II) sulfate pentahydrate and THPTA was mixed first, followed by the addition of sodium ascorbate to be final molar ratio of 1:2:2. A 37-fold molar excess of Cu-THPTA complex was added to tube 1 and water and DMSO were added to get the final reaction mixture with 50% DMSO. Click reaction mixture was mixed thoroughly and flushed with inert gas (N_2_) for 1 minute. Reaction mixture was then incubated at room temperature for ∼ 16 h. Click reaction product was purified by reverse phase high-performance liquid chromatography (HPLC) using a C18 column. HPLC method used for oligo purification (Buffer A-5% acetonitrile, 4.25% Triethylamine acetate (TEAA) in water; Buffer B-100% acetonitrile (ACN). The program was set with a flow rate of 5 mL/min for 150 minutes, and a gradient of ACN increasing from 0-80%.

### Annealing reaction

FPLC purified single stranded oligo conjugated to VHL (oligo-VHL) ligand and its reverse complement oligo were dissolved in ultra-pure water. Single stranded oligo-VHL and single stranded reverse complement oligos were mixed 1:1 molar ratio (final concentrations of TRAFTACs were set to 25 µM) in 1X annealing buffer (10 mM Tris, pH 7.5, 50 mM NaCl and 1 mM EDTA) and incubated for 10 minutes in a water bath at 95 degrees Celsius. Then, the hot-plate was turned off and the samples were left in the water bath and let cool down to room temperature over 2 h. Double stranded oligoTRAFTACs were mixed by gently vortexing and aliquoted and stored at -20 degrees Celsius.

### Western blotting

Protein concentration in all the cell lysates were measured by BCA protein assay kit and equal amounts from each lysate were mixed with 4X loading dye and boiled for 5 minutes followed by 2 minutes centrifugation prior to loading into SDS-PAGE gel. Next proteins on the SDS-PAGE gel were transferred to a PVDF membrane by western blotting and the membrane was blocked with 5% milk in TBST (0.05%Tween 20) for 1 h. Primary antibodies (all Cell Signaling antibodies were diluted 1:1000, c-Myc 1:150) were prepared in TBST with 5% milk and membranes were incubated overnight at 4°C. On the following day, membrane was washes for 15 minutes (Incubate for three times, 5 minutes each) and appropriate secondary antibodies (1:5000) were prepared in TBST with 5% milk and incubated with the membrane for 1h at room temperature (RT). Membrane was washed for 30 minutes with TBST (incubate for three times, 10 minutes each) prior to imaging.

### EMSA

Click reaction mixtures and unreacted alkyne-modified oligonucleotides were separated in a 1.2 % agarose gel for 1 h at constant 120 mV and DNA bands were captured by the ChemiDoc system (BioRad) using SYBR safe mode.

### Biotin pull down

Cells (HEK293T or HeLa) were grown in three T-175 flasks for 2 days. When cells reach >90% confluency, cells were harvested and washed once with 1X PBS. Cells were pooled together and resuspended in 1.5 mL immunoprecipitation buffer (25 mM Tris pH 7.4, 150 mM NaCl, 0.4% NP40, 5% glycerol, 1X protease inhibitor cocktail from Roche and 1 mM of PMSF). Cells were then incubated for 30 minutes on ice prior to centrifugation at high speed (15,000 rpm) for 20 minutes. Equal amounts/volumes (∼1 mg) of clarified lysate were transferred to individual tubes and incubated with biotinylated double stranded oligonucleotides for 2 h at RT. Pre-washed 30 µl of streptavidin agarose beads were transferred to each tube and incubated overnight at 4°C. Beads were then washed with 1X TBS for 15 minutes (three times, 5-minute incubation each time). Bound proteins were eluted with 2X loading buffer by boiling for 8 minutes. Boiled samples were centrifuged at high speed for 5 minutes and supernatant was loaded onto SDS-PAGE gel followed by western blotting.

### Immunoprecipitation and ubiquitination assay

A HA-tagged ubiquitin plasmid (4 µg) was transfected into HEK293T cells overexpressing brachyury-GFP in a 10 cm dish. On the following day, transfected cells were split into three 10 cm cell culture dishes and incubated for 24 h prior to transfection of oligoTRAFTACs. Epoxomicin (1 µM) was preincubated with cells for 1 h and oligoTRAFTACs (mock, OT3 and OT17) were then transfected using RNAi-MAX in 5 mL/dish of transfection medium. After 12 h post transfection, cells were harvested and lysed using immunoprecipitation buffer (25 mM Tris pH 7.4, 150 mM NaCl, 0.4% NP40, 5% glycerol, 1X protease inhibitor cocktail from Roche and 1 mM of PMSF). Approximately 1.5 mg of lysate from each sample was incubated with brachyury antibody at 4°C for 4 h. Protein A agarose beads (30µl) were added to antibody, lysates mixture and rock at 4°C for ∼ 18 h. Beads were washed with 1X TBS for three times with 5-minute incubation during each wash. Immunoprecipitated proteins were eluted by boiling agarose beads in 2X loading buffer (containing 10% ß-ME) for 8 minutes and centrifuged at high speed for 5 minutes prior to the loading into SDS-PAGE gel followed by western blot analysis.

### Cell viability assay

Cells were split and subjected to oligoTRAFTAC transfection as described in ‘‘oligoTRAFTAC transfection” method section. Following the transfection, cells were incubated for the appropriate number of days before recording the luminescence reading from the plate reader. CellTiter-Glo reagent was prepared according to the manufactures recommendation and mixed with cell culture medium with 1:1 ratio. CellTiter-Glo reagent (100 µl) was added to each well and incubated for 10 minutes before taking the reading.

### Zebrafish Husbandry and Microinjection

Tüpfel-longfin zebrafish were raised according to standard protocols approved by the Institutional Animal Care and Use Committee. Experiments were performed before sex determination in zebrafish (*40*). Embryos were injected at the one cell stage with 180 picoliters of a 25 µM oligoTRAFTAC solution. They were raised for 48 hours at 28.6 °C and then scored for the presence of tail defects.

## Supporting Figures

**Figure S1.**
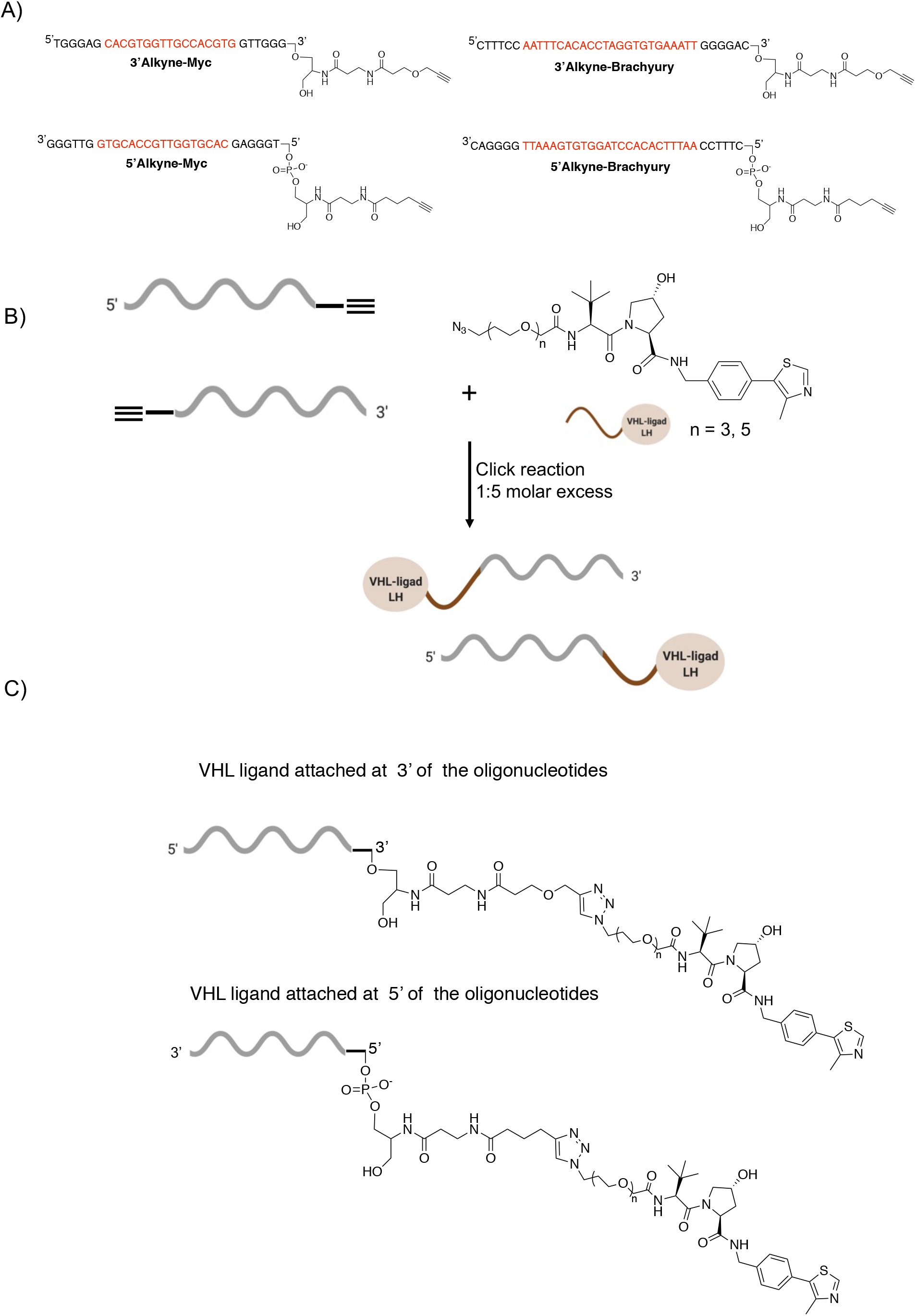
Oligonucleotide synthesis and click reaction. A) Oligonucleotides were custom synthesized with a terminal alkyne either at 3’ or 5’ end of the oligo. Oligonucleotide sequences for both 3’ and 5’ alkyne targeting c-Myc (left panel) and brachyury (right panel). B) Chemical structure of azido-VHL ligand and click reaction conditions. C) Chemical structures of 3’ and 5’ modified oligonucleotides after the click reaction with azido-VHL ligand.

**Figure S2.**
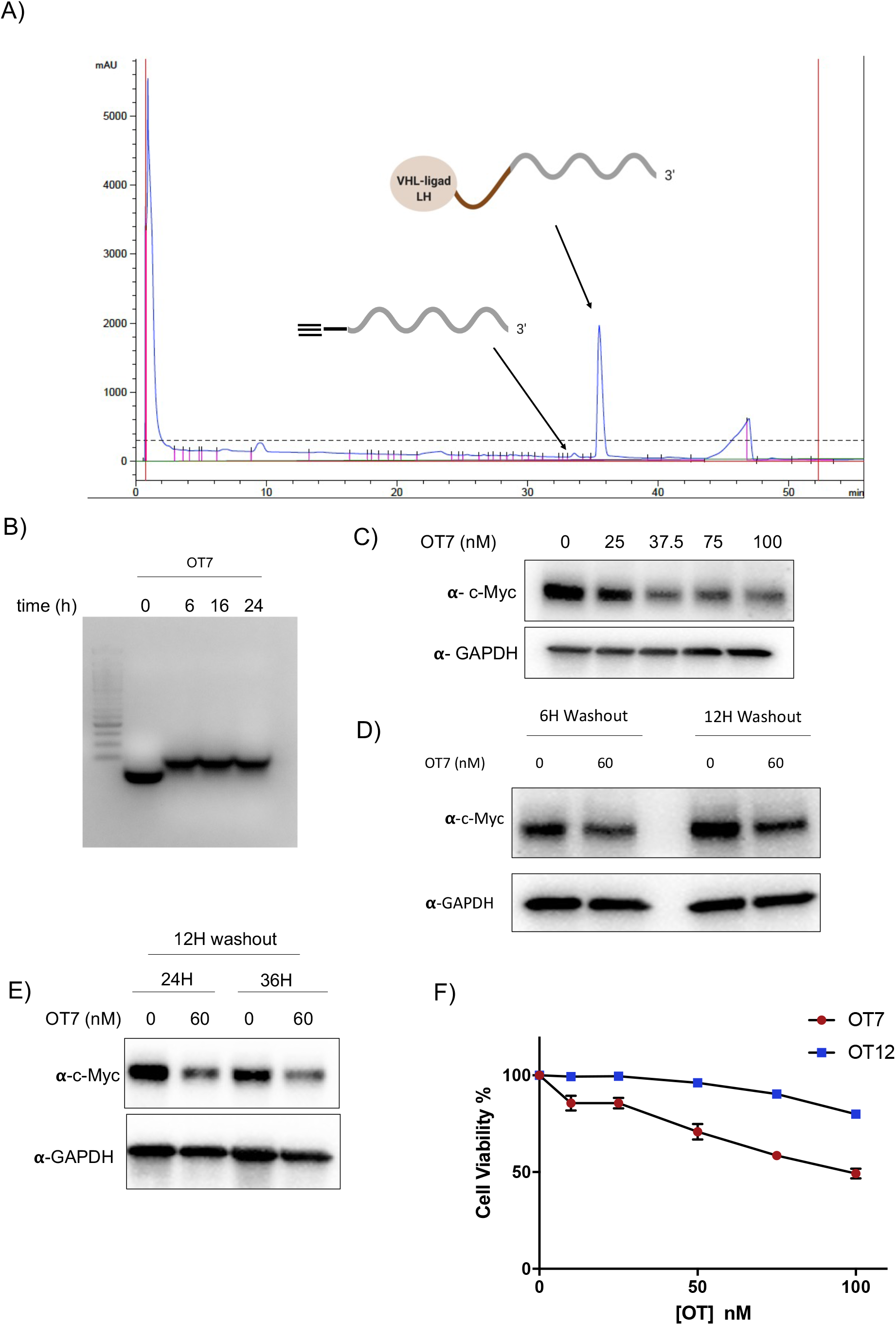
OligoTRAFTAC mediated c-Myc degradation. A) HPLC trace for crude click reaction mixture of OT7. B) Gel shift assay data for the click reaction. Before and after click reaction, oligonucleotides were separated from a 1.2 % agarose gel for 1 h at constant 120 mV. VHL ligand reacted oligonucleotides were shifted compared to the unreacted oligo. C) Varying concentrations of OT7 were transfected into HEK293T cells and lysed after 20 h. Cell lysates were separated and transferred to a PVDF membrane followed by immunoblotting with antibodies against c-Myc and GAPDH. D) After 60 nM of OT7 transfection, cell culture media was replaced with fresh medium and continued the incubation for total of 20 h prior to lysis. Lysates were probed for c-Myc and GAPDH. E) Similarly, HEK293T cells were subjected to a washout experiment 12 h post-transfection and lysed after 24 h and 36 h. F) HeLa cells were seeded into 96-well plates transfected increasing concentrations of OT7 and OT12. After 48 h, cell viability was monitored using CellTiter-Glo reagent.

**Figure S3.**
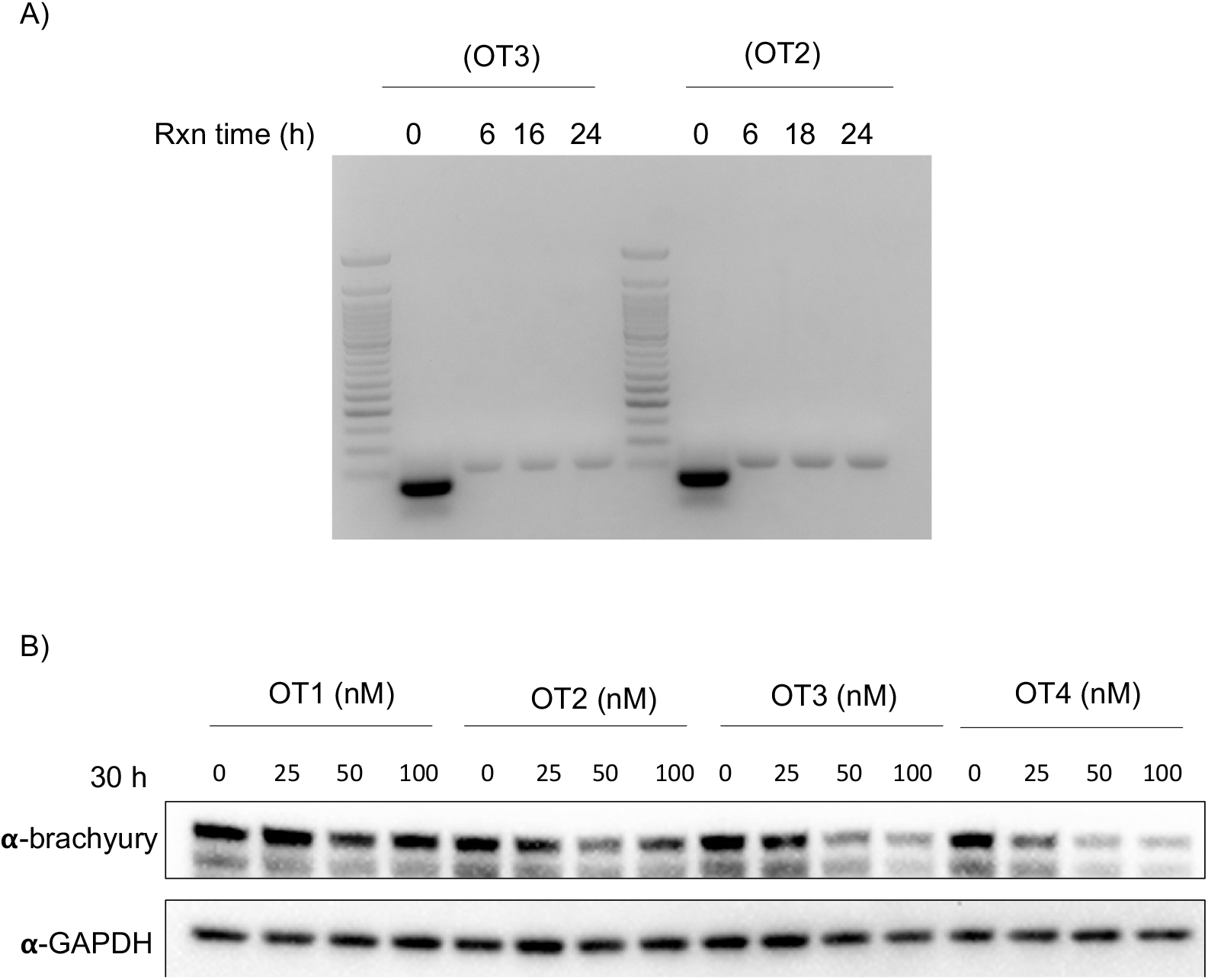
EMSA and brachyury-GFP degradation data. A) Click reaction mixtures with or without VHL ligand were loaded on to a 1.2% agarose gel and separated over 1 h at constant 120 mV. B) OT1 through 4 were transfected into HEK293T cells overexpressing brachyury-GFP and lysed after 30 h. Cell lysates were subjected to SDS-PAGE and western blotting followed by probing with antibodies against brachyury and GAPDH.

**Figure S4.**
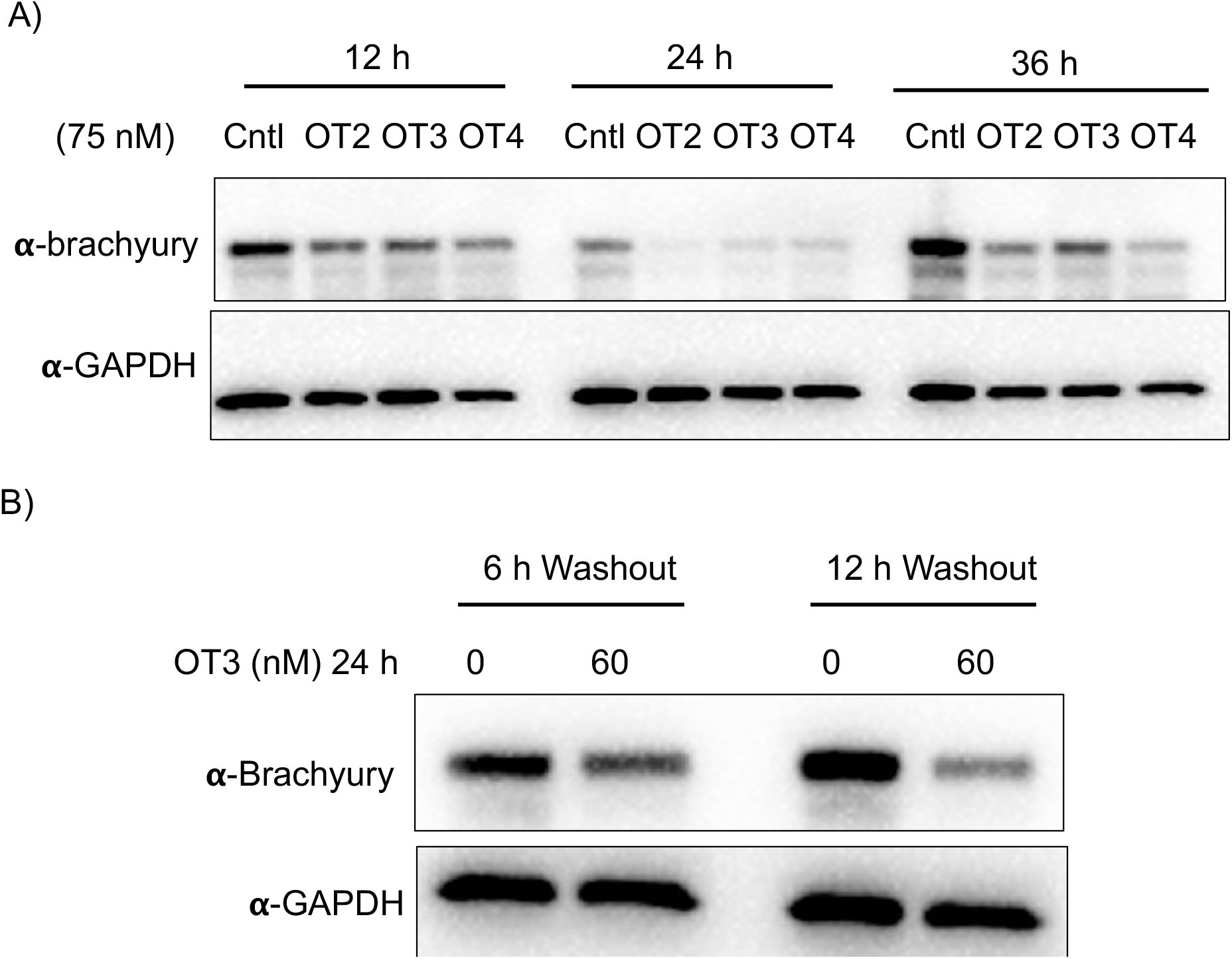
Time course and washout experiments for brachyury targeting OTs. A) OT2 through 4 were transfected at 75 nM into HEK293T cells overexpressing brachyury-GFP and lysed after 12h, 24 h, and 36 h. Lysates were probed for brachyury and GAPDH. B) OT3 was transfected into HEK293T cells and washed out after 6 h and 12 h. Cell lysates were probed with antibodies against brachyury and GAPDH.

**Figure S5.**
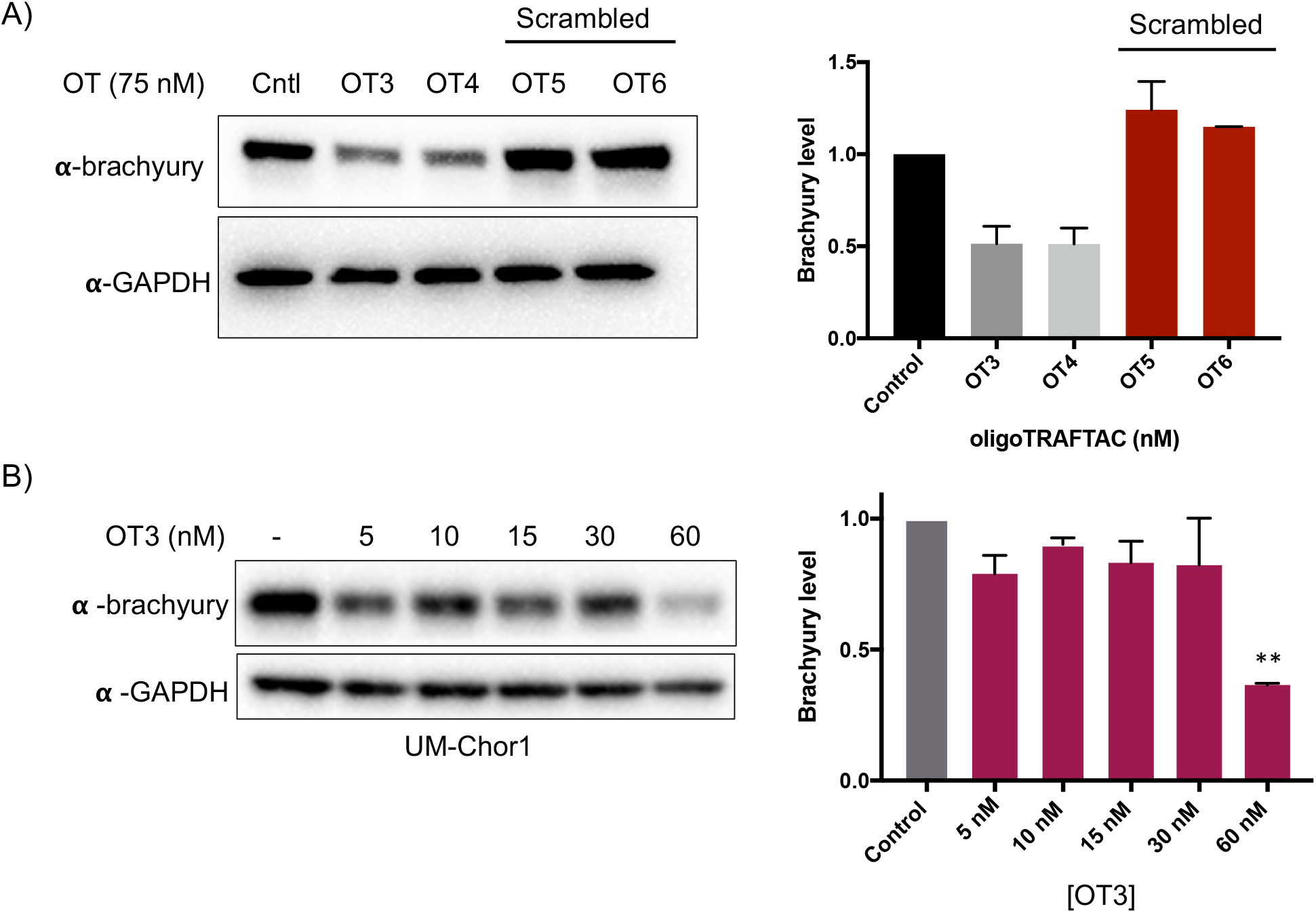
Brachyury degradation by oligoTRAFTACs in HEK293T and UM-Chor1 cells. A) Brachyury-GFP degradation by OTs is sequence dependent. OT3, OT4, and their scrambled OTs (OT5 and OT6) were transfected and lysed after 20 h. Cell lysates were subjected to SDS-PAGE and western blotting. Blots were probed with brachyury and GAPDH antibodies. Quantitation of western blot bands is shown on the right. B) Increasing concentration of OT3 were transfected into UM-Chor1 cells and degradation was evaluated after 24 h.

### General chemistry methods

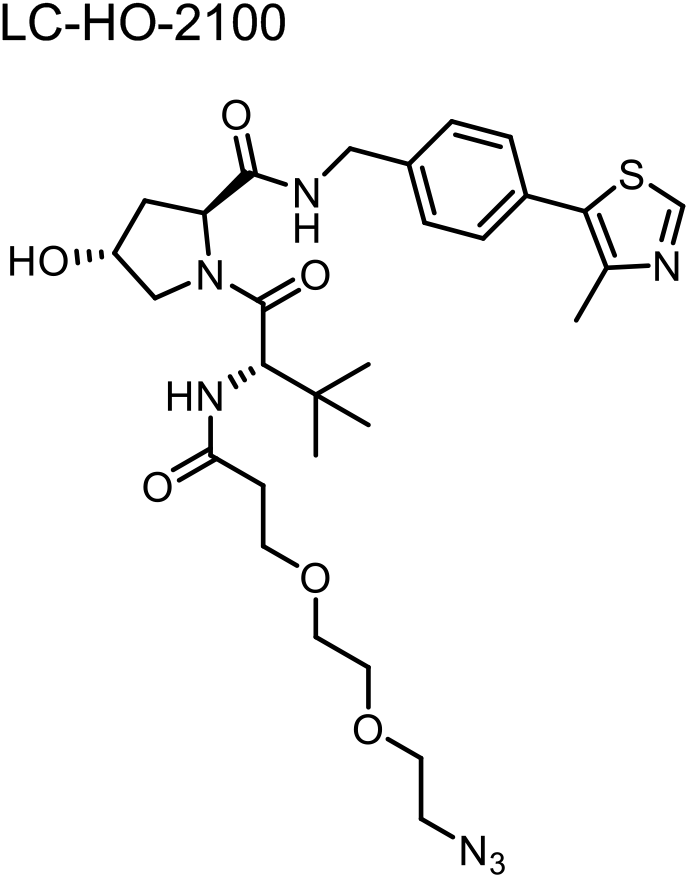

To a solution of (2S,4R)-1-[(2S)-2-amino-3,3-dimethylbutanoyl]-4-hydroxy-N-[[4-(4-methylthiazol-5-yl)phenyl]methyl]pyrrolidine-2-carboxamide hydrochloride (1.0 eq., 20 mg) in DMF (1 mL) was added (2,5-dioxopyrrolidin-1-yl) 3-[2-(2-azidoethoxy)ethoxy]propanoate (1.2 eq.), and TEA (5.0 eq.). The mixture was stirred for 1 hour at room temperature. Upon completion, the mixture was diluted with H_2_O (5 mL), extracted with EtOAc (3×5 mL), dried over Na_2_SO_4_, concentrated under vacuum and purified by prep-TLC. The title compound (23 mg, 71% yield) was obtained as a colorless oil.

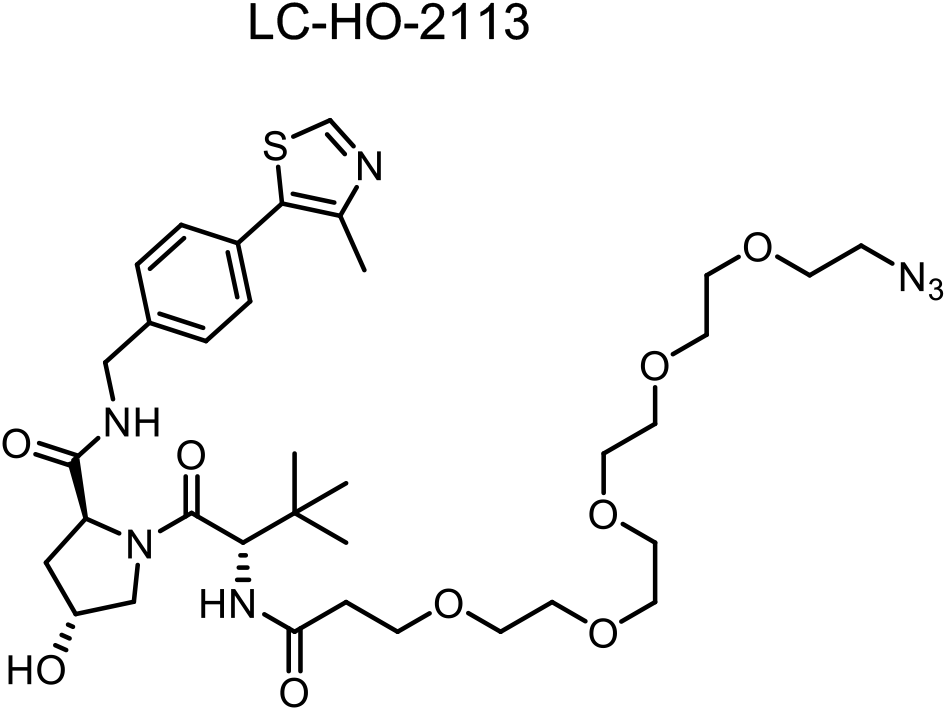

To a solution of (2S,4R)-1-[(2S)-2-amino-3,3-dimethylbutanoyl]-4-hydroxy-N-[[4-(4-methylthiazol-5-yl)phenyl]methyl]pyrrolidine-2-carboxamide hydrochloride (1.0 eq., 20 mg) in DMF (1 mL) was added(2,5-dioxopyrrolidin-1-yl)3-[2-[2-[2-[2-(2-azidoethoxy)ethoxy]ethoxy]ethoxy]ethoxy]propanoate (1.2 eq.), and TEA (5.0 eq.). The mixture was stirred for 1 hour at room temperature. Upon completion, the mixture was diluted with H_2_O (5 mL), extracted with EtOAc (3×5 mL), dried over Na_2_SO_4_, concentrated under vacuum and purified by prep-TLC. The title compound (19 mg, 57% yield) was obtained as a colorless oil.

Chemical shifts are reported in δ ppm referenced to an internal CDCl_3_ (δ 7.26 ppm) for ^**1**^**H NMR**, CDCl_3_ (δ 77.00) for ^**13**^**C NMR**.

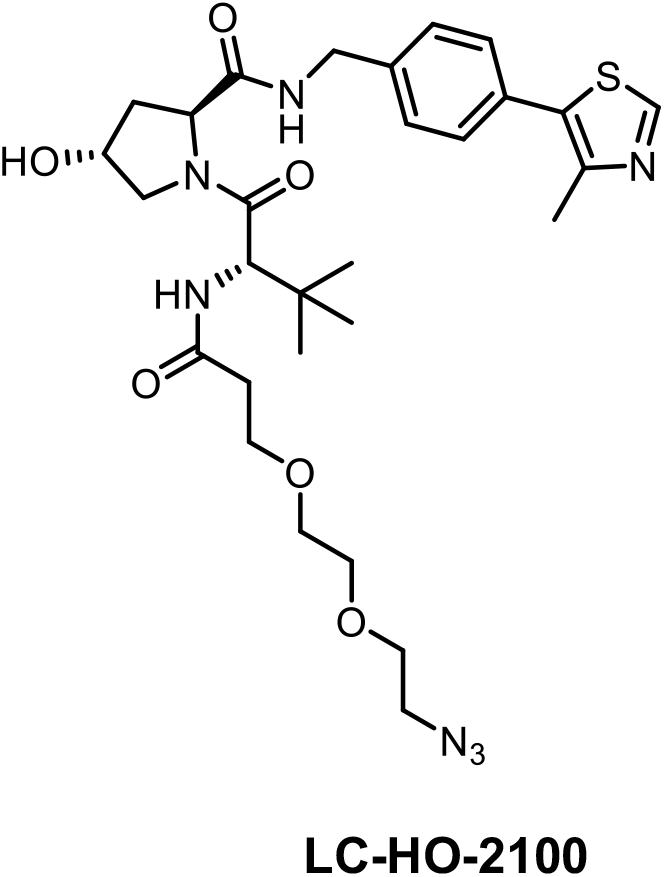

^**1**^**H NMR** (600 MHz, CDCl_3_) δ 8.67 (s, 1H), 7.42 (t, *J* = 6.0 Hz, 1H), 7.36 – 7.32 (m, 4H), 6.99 (d, *J* = 8.4 Hz, 1H), 4.70 – 4.67 (m, 1H), 4.54 – 4.51 (m, 1H), 4.47 – 4.46 (m, 1H), 4.11 (d, *J* = 11.4 Hz, 1H), 3.73 – 3.69 (m, 3H), 3.64 (s, 3H), 3.65 – 3.62 (m, 3H), 3.59 (dd, *J* = 10.8, 3.6 Hz, 1H), 3.37 – 3.35 (m, 2H), 2.50 (s, 3H), 2.53 – 2.47 (m, 3H), 2.13 – 2.09 (m, 3H), 0.93 (s, 9H).

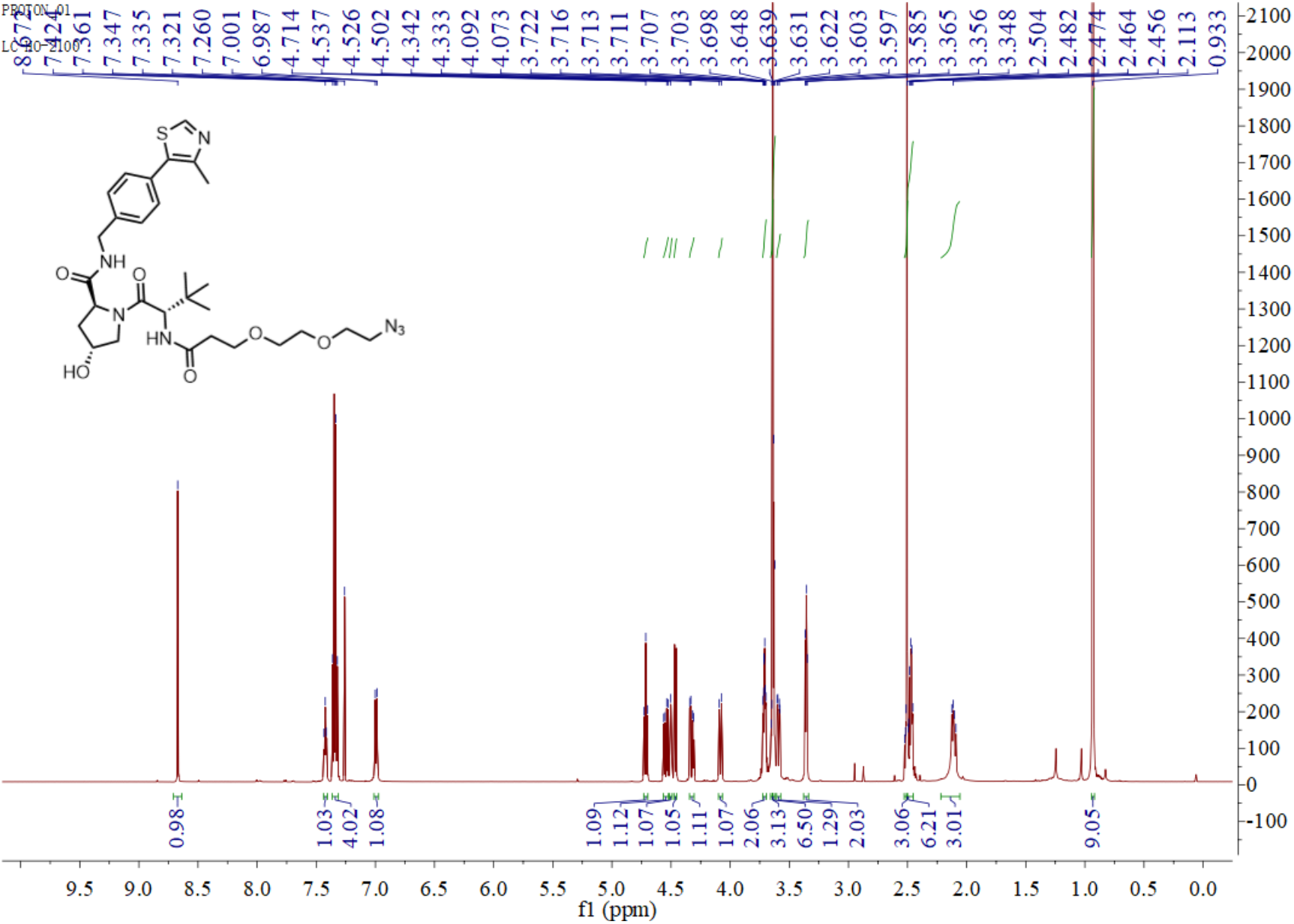

^**13**^**C NMR** (150 MHz, CDCl_3_) δ 172.07, 171.71, 170.76, 150.28, 148.40, 138.10, 131.55, 130.87, 129.45, 128.06, 70.45, 70.42, 70.02, 69.97, 67.13, 58.37, 57.67, 56.58, 50.52, 43.17, 36.65, 35.82, 34.78, 26.35, 16.02.

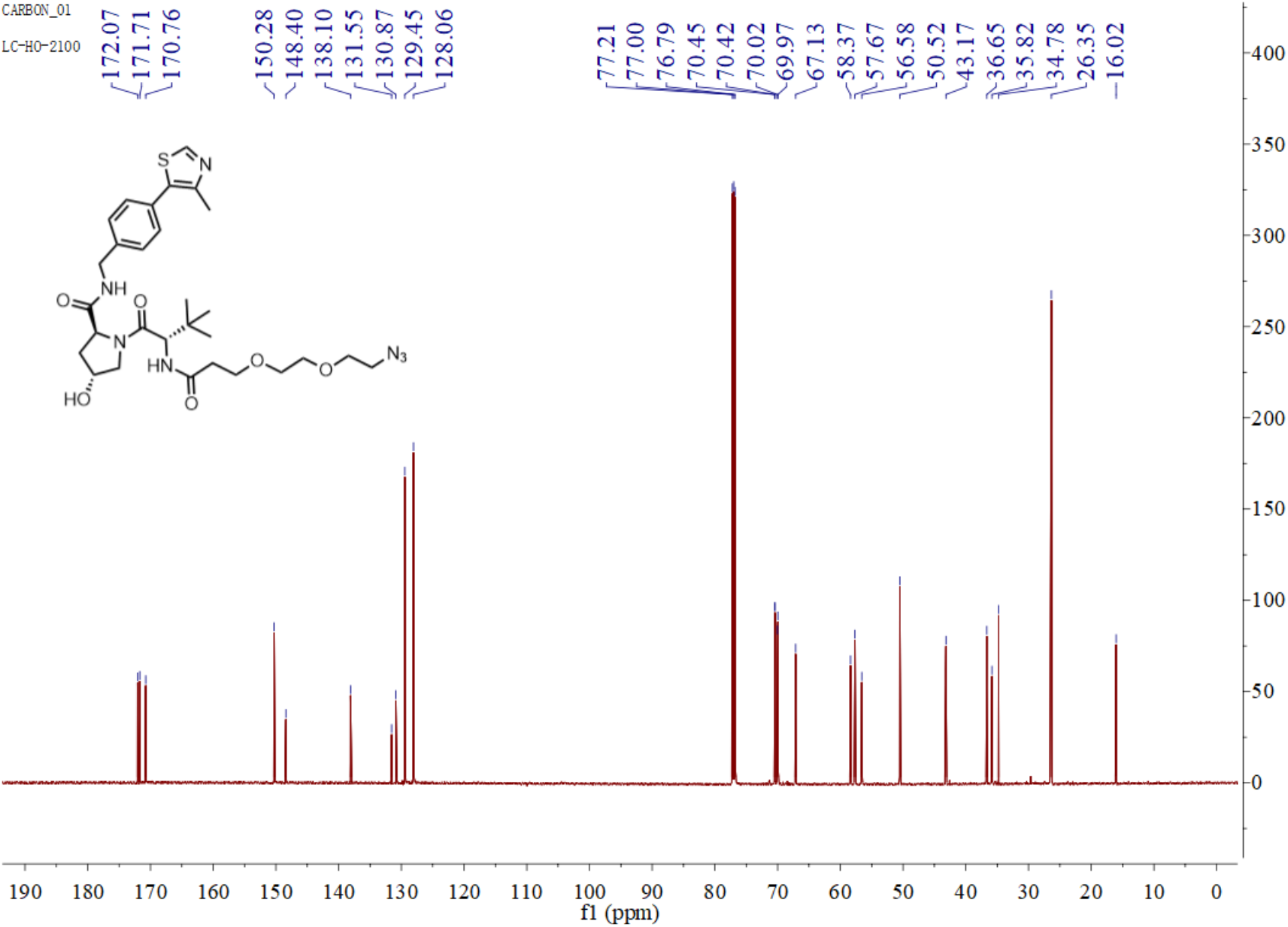

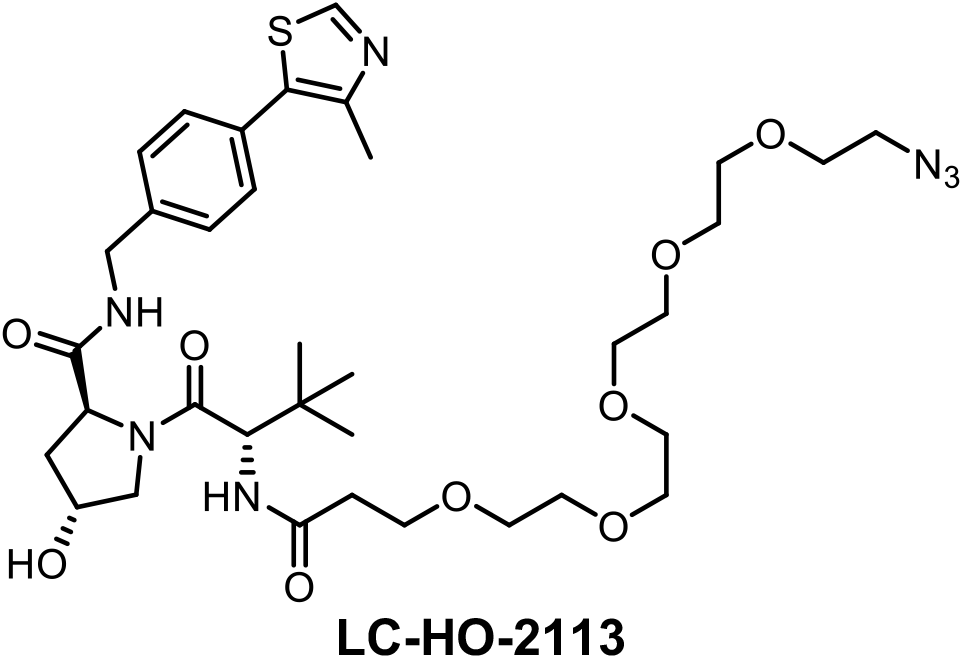

^**1**^**H NMR** (600 MHz, CDCl_3_) δ 8.66 (s, 1H), 7.47 (s, 1H), 7.34 – 7.33 (m, 4H), 7.04 –7.03 (m, 1H), 4.71 – 4.67 (m, 1H), 4.55 – 4.50 (m, 1H), 4.49 – 4.44 (m, 2H), 4.33 – 4.29 (m, 1H), 4.05 (d, *J* = 10.8 Hz, 1H), 3.69– 3.66 (m, 3H), 3.65 – 3.58 (m, 17H), 3.36 – 3.35 (m, 2H), 2.49 – 2.48 (m, 4H), 2.48 – 2.38 (m, 4H), 2.13 – 2.07 (m, 1H), 0.92 (s, 9H).

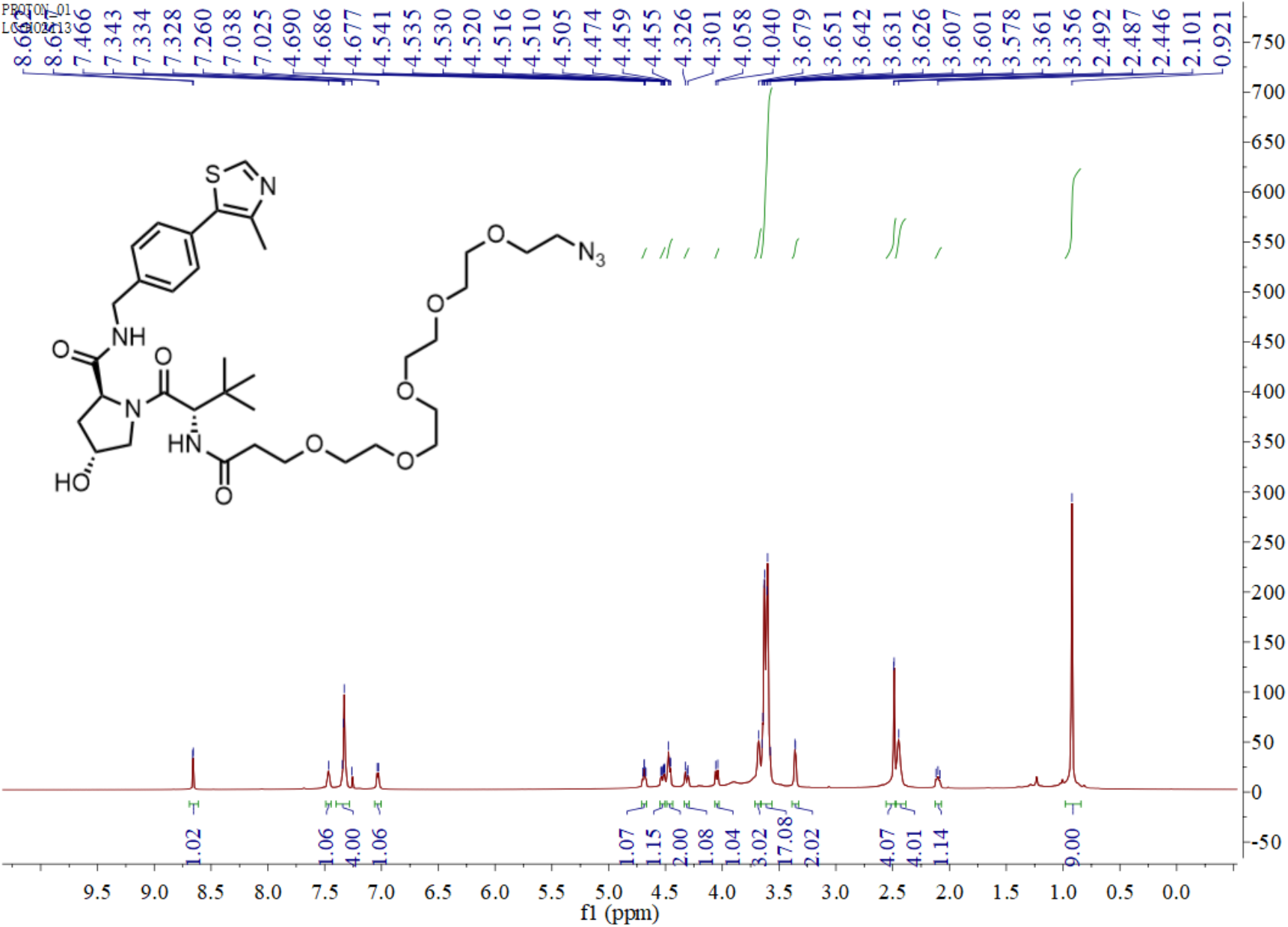

^**13**^**C NMR** (150 MHz, CDCl_3_) δ 172.01, 171.59, 170.91, 150.25, 148.34, 138.16, 131.55, 130.76, 129.39, 127.99, 70.59, 70.54, 70.50, 70.44, 70.41, 70.37, 70.32, 69.97, 69.95, 67.04, 58.45, 57.65, 56.62, 50.59, 43.08, 36.61, 36.03, 34.87, 26.33, 15.99.

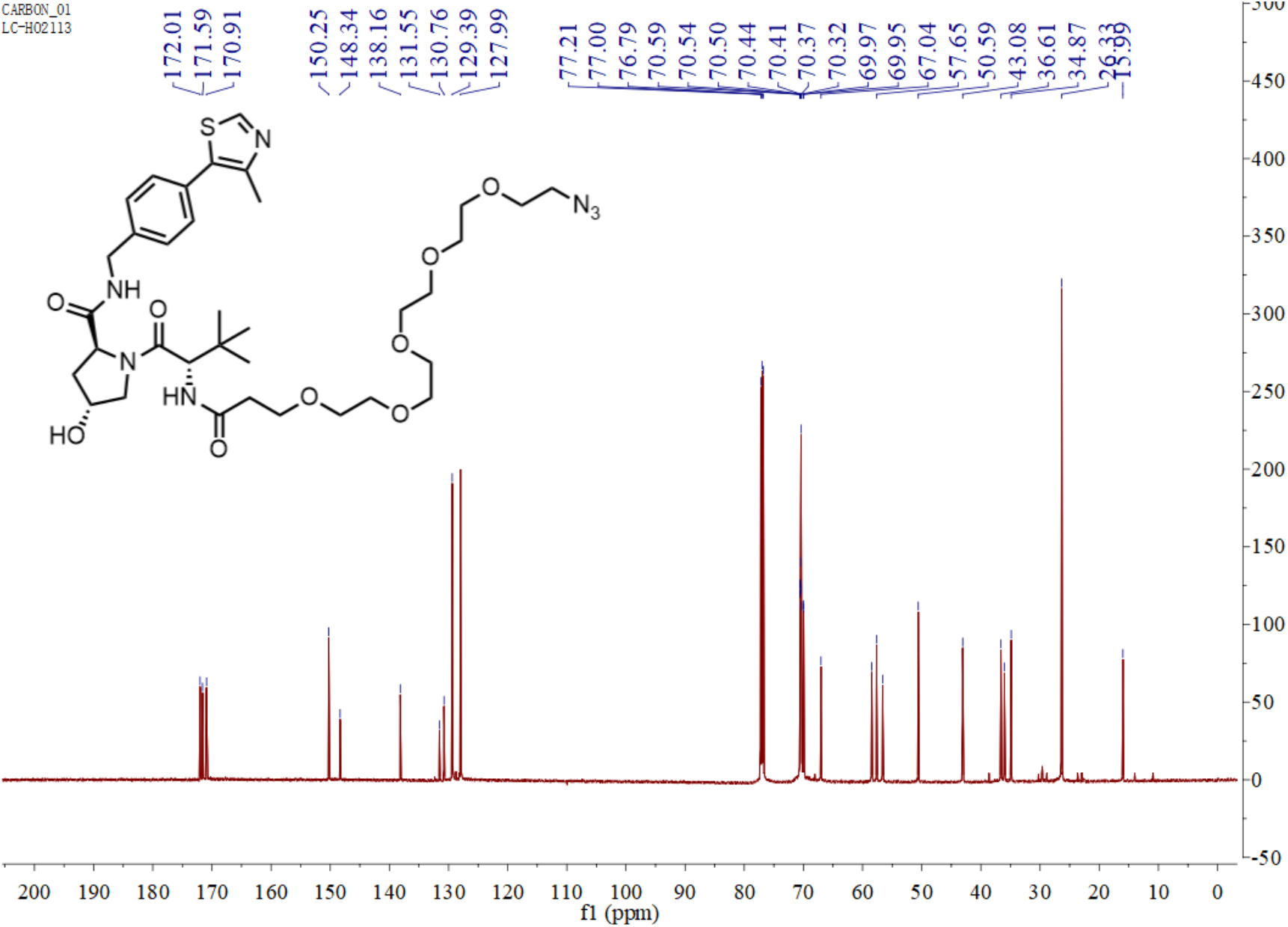

